# Control of cell proliferation by memories of mitosis

**DOI:** 10.1101/2022.11.14.515741

**Authors:** Franz Meitinger, Robert L. Davis, Mallory B. Martinez, Andrew K. Shiau, Karen Oegema, Arshad Desai

**Affiliations:** Department of Cell and Developmental Biology, School of Biological Sciences, University of California San Diego, La Jolla, CA 92093, USA; Department of Cellular and Molecular Medicine, University of California San Diego, La Jolla, California 92093, USA; Small Molecule Discovery Program, Ludwig Institute for Cancer Research, La Jolla, California 92093, USA; Ludwig Institute for Cancer Research, La Jolla, California 92093, USA

## Abstract

Mitotic duration is tightly constrained, with extended mitotic duration being a characteristic of potentially problematic cells prone to chromosome missegregation and genomic instability. We show that memories of mitotic duration are integrated by a p53-based mitotic stopwatch pathway to exert tight control over proliferation. The stopwatch halts proliferation of the products of a single significantly extended mitosis or of successive modestly extended mitoses. Time in mitosis is monitored via mitotic kinase-regulated assembly of stopwatch complexes that are transmitted to daughter cells. The stopwatch is inactivated in p53-mutant cancers, as well as in a significant proportion of p53-wildtype cancers, consistent with classification of stopwatch complex subunits as tumor suppressors. Stopwatch status additionally influences efficacy of anti-mitotic agents currently used or in development for cancer therapy.

**One-Sentence Summary:** Time spent in mitosis is carefully monitored to halt the proliferation of potentially dangerous cells in a population.

## Introduction

Mitosis is a complex event that is executed in a narrow temporal window of the cell cycle. Mitotic duration is tightly constrained (Araujo et al., 2016), and prolonged mitotic duration is a sign of problems that can lead to chromosome missegregation, a trigger event for genomic instability (Shoshani et al., 2021; Umbreit et al., 2020). Monitoring mitotic duration is therefore an effective means to identify potentially problematic cells in a proliferative population. Prior work has shown that when a threshold mitotic duration is surpassed, daughter cells arrest in G1 in a p53-dependent manner (Uetake and Sluder, 2010). In addition to p53 and its transcriptional target p21, two additional factors–the scaffold protein 53BP1 and the deubiquitinase USP28–are required for daughter cell arrest following extended mitotic duration (**Fig. 1A,B; Fig. S1A,B;** (Fong et al., 2016; Lambrus et al., 2016; Meitinger et al., 2016)). As daughter cell arrest is independent of the cause of the extended mitotic duration (Uetake and Sluder, 2010), the monitoring mechanism functions akin to a stopwatch. When spindle assembly is prolonged by genetically-engineered centriole removal in the early mouse embryo or the developing nervous system, the observed cell death, embryonic arrest, and microcephaly phenotypes are ameliorated by removal of stopwatch components, indicating that this pathway is operational during development (Bazzi and Anderson, 2014; Insolera et al., 2014; Marjanovic et al., 2015; Phan and Holland, 2021; Phan et al., 2021; Xiao et al., 2021). However, how the stopwatch measures mitotic duration to control daughter cell fate and the physiological significance of stopwatch-mediated control of cell proliferation remain unknown.

### A memory of mitotic duration is transmitted to control daughter cell G1

Given the sharp transition in daughter cell proliferation versus arrest as a function of mother cell mitotic duration (**Fig. 1B**), we first addressed if the stopwatch involves a threshold event in mitosis or if a memory of mitotic duration is transmitted to daughter cells, where the threshold decision is made. Given that the output of the stopwatch pathway, p21, is necessary for daughter cell G1 arrest (**Fig. 1B**), we *in situ-tagged* p21 with mNeonGreen (mNG) to distinguish between these possibilities. In the p21-mNG line, the threshold mother cell mitotic duration for daughter cell arrest was ~50% longer than in the parental line (~150 min vs ~100 min), likely due to the fluorescent tag (*not shown*). We correlated mother cell mitotic duration with p21-mNG expression in daughter cells to determine whether p21 is only elevated in the daughters of mothers whose mitotic duration surpasses the arrest threshold, or whether increasing mitotic durations are encoded in progressively higher p21 levels in G1 that trigger arrest when a threshold is surpassed. Following mitoses with a normal duration of ~30 min, p21 expression was low in most cells (**Fig. 1C,D**). Extending mitotic duration to sub-threshold times between 60 and 150 minutes progressively increased p21 levels (**Fig. 1C,D; Fig. S1C**). Notably, for all cells, p21-mNG signal was only observed after transit into G1 (**Fig. 1C; Fig. S1C**). For sub-threshold mitotic durations, p21-mNG signal was detected in G1 and then abruptly dropped, likely due to ubiquitin-dependent degradation in early S-phase (Barr et al., 2017); p21-mNG was then detected again in G2 prior to the next M-phase (**Fig. 1C**). Above the ~150 min threshold, high p21-mNG signal in G1 persisted and no subsequent M-phase was observed (**Fig. 1C**). Using the drop in p21 signal as a marker for the G1-S transition, G1 duration, like p21-mNG expression, progressively increased for subthreshold mitotic durations between 30 and 150 minutes (**Fig. 1D**); a similar progressive increase in G1 duration, but not in S+G2 duration, was also observed using a cell cycle phase sensor in a cell line without tagged p21 (**Fig. S1D,E**). These results suggest that the stopwatch transmits a memory of mitotic duration in an analog fashion to daughter cells, leading to a progressive increase in p21 levels with increased mitotic time. The fact that this progressive increase is converted into a sharp threshold in mother cell mitotic time for G1 arrest is consistent with p21, a stoichiometric inhibitor of G1 Cdk complexes, exhibiting ultrasensitivity (Ferrell and Ha, 2014; Yang et al., 2017).

### Memory of extended mitotic duration is integrated across cell cycles to control cell proliferation

The results above suggest that a memory of mitotic duration is transmitted to immediately produced daughter cells. We next assessed if a memory of mitotic duration can be transmitted in a multigenerational manner across successive cell cycles. For this purpose, we developed a means to extend average mitotic duration from ~30 min to ~60 min, well below the ~100 min threshold for immediate daughter cell G1 arrest (**Fig. 1E**). We achieved this through inducible inactivation of a mechanism that promotes, but is not essential for, anaphase onset (*iComet*Δ; **Fig. 1E, Fig. S1F,G**; (Corbett, 2017; Kim et al., 2018)). Notably, *iComet*Δ, while delaying anaphase onset, does not perturb spindle assembly or chromosome alignment and segregation (**Fig. S1H**). Despite mother cells transiting mitosis in comparable time intervals and without visible segregation errors, the frequency of daughter cell arrest following sub-threshold mitotic durations of 60-90 minutes increased progressively between days 2 and 4 after induction of *Comet* deletion, as cells experienced successive sub-threshold extended mitoses (**Fig. 1E,F; Fig. S1G**). Notably, daughter cell arrest was greatly reduced for *iComet*Δ in cell lines lacking the essential stopwatch components USP28 or p53 (**Fig. 1F; Fig. S1I**). Thus, a stopwatch-dependent memory of subthreshold mitotic extension is transmitted and integrated across sequential cell cycles to control cell fate. An orthogonal perturbation that leads to a sub-threshold extension of mitosis by delaying spindle assembly, coupled with imaging of p21-mNG, provided additional support for this conclusion (**Fig. S2**).

### Stopwatch complexes transmit memory of mitotic duration

To understand how memory of extended mitosis controls G1 progression, we conducted biochemical analysis of 53BP1 and USP28, components required for stopwatch function, in extracts prepared from mitotically arrested and asynchronously cycling (primarily interphase) cells. While USP28 was soluble in both, 53BP1 was soluble in mitotic but largely insoluble in interphase extracts (**Fig. 2A,B; Fig. S3A**); low solubility of 53BP1 in interphase is consistent with its chromatin association (Fernandez-Vidal et al., 2017; Zimmermann and de Lange, 2014). Immunoprecipitation of 53BP1 from different cell cycle states revealed robust coimmunoprecipitation of 53BP1, USP28 and p53 specifically from mitotic extracts (**Fig. 2C; Fig. S3B**). Neither an interaction between 53BP1 and USP28 nor increased 53BP1 solubility was detected following DNA damage, despite robust p53 elevation, indicating that complex formation in mitosis is not a secondary consequence of DNA damage (**Fig. 2D; Fig. S3A**). Immunoprecipitation of 53BP1 from cells lacking USP28 or p53 revealed that 53BP1 independently interacts with the other two complex components (**Fig. 2E**), which is consistent with the C-terminal tandem BRCT domain of 53BP1 interfacing with p53 (Derbyshire et al., 2002; Joo et al., 2002), and a point mutation (G1560K) in the Tudor domain of 53BP1 disrupting interaction with USP28 (Cuella-Martin et al., 2021). Introducing the G1560K mutation using base editing selectively disrupted the interaction of 53BP1 with USP28 in mitotically arrested cells (**Fig. 2F, Fig. S3C**), and inhibited stopwatch function to a similar extent as *TP53BP1* knockout (**Fig. 2G**). Collectively, these results indicate that 53BP1 employs distinct interfaces to bind USP28 and p53 to scaffold the mitosis-specific formation of stopwatch complexes that transmit the memory of mitotic duration to control G1 progression.

To address how mitotic stopwatch complexes form specifically in mitosis, we performed a focused screen with chemical inhibitors targeting 7 mitotic and DNA damage kinases (**Fig. 3A**); targeted kinases were limited by the need for cells to enter and prolong mitosis in presence of a reversible spindle assembly inhibitor (**Fig. 3A**), which excluded CDK1 and Aurora B. This screen identified the mitotic kinase PLK1 as being essential for the mitotic stopwatch (**Fig. 3A**). By contrast, the DNA damage response kinases ATM, ATR, DNA-PK, CHK1 and CHK2 did not contribute to stopwatch function (**Fig. 3A; Fig. S4A-C**), and inhibition of Aurora A had only a minor effect (**Fig. S4D**). A chemically distinct PLK1 inhibitor and introduction of a mutation conferring partial PLK1 inhibitor resistance, confirmed the specific requirement for PLK1 activity (**Fig. S4E,F**). Notably, PLK1 activity was essential for the independent 53BP1–USP28 and 53BP1–p53 interactions but was not required for the mitosis-specific increase in 53BP1 solubility (**Fig. 3B**). Thus, formation of mitotic stopwatch complexes is triggered by PLK1 kinase activity, acting in conjunction with mitosis-specific solubility of 53BP1 (**Fig. 3D**).

To transmit a memory of mitotic duration, stopwatch complexes must persist into G1 to control p21 expression. Consistent with this expectation, 6 hours after release from mitotic arrest into G1 by inhibitor washout, 53BP1 remained soluble and stopwatch complexes were detected at levels comparable to mitotic arrest (**Fig. 3C**). Notably, p21 was only induced after the transition to G1; in addition, more p53 was detected in G1 stopwatch complexes, and p53 levels were higher overall at this cell cycle stage **(Fig. 3C)**. These results suggest a model in which there is a mitotic time and PLK1-dependent formation of stopwatch complexes containing USP28 and 53BP1 that recruit available p53. Within stopwatch complexes, 53BP1 positions p53 in proximity to USP28, leading to p53 deubiquitination and stabilization. The soluble 53BP1—USP28 complexes formed in a PLK1 activity-dependent manner persist into G1, where they likely continue to deubiquitinate and stabilize p53, which in turn induces p21 synthesis (**Fig. 3D**).

### Stopwatch inactivation in p53-wildtype cancers

Difficulty in executing events such as spindle assembly or chromosome-spindle attachment extends mitosis via activation of a checkpoint and is associated with elevated rates of chromosome missegregation (Cimini et al., 2001), which in turn lead to aneuploidy and genomic instability. The mitotic stopwatch monitors mitotic duration and halts the proliferation of cells that experience either one highly delayed mitosis or successive moderately delayed mitoses, making it well suited to act as a fidelity filter that suppresses proliferation of potentially dangerous cells in a population. Consistent with this notion, the genes encoding the three stopwatch complex subunits are ranked among the top 100 tumor suppressor genes ((Davoli et al., 2013); **Fig. S4G,H**). Classification of *USP28* as a tumor suppressor gene is particularly significant as, unlike *TP53BP1 and TP53*, it is not an integral component of the DNA damage response (Knobel et al., 2014). As stopwatch function relies on p53 (Uetake and Sluder, 2010), the stopwatch is predicted to be inactive in the ~50% of human cancers that harbor p53 mutations. This expectation was confirmed in three p53-mutant cancer-derived cell lines (**Fig. 4A,B**; **Fig. S5A,B**). A more interesting question, relevant to addressing the physiological function of the stopwatch, is its status in the ~50% of cancers that express wildtype p53 (Khoo et al., 2014; Olivier et al., 2010). To determine the status of the stopwatch in p53-wildtype cancers, we surveyed 15 cell lines from pediatric and adult cancers that were annotated as expressing wildtype p53 (**Fig. 4A**). For all tested cell lines, expression of functional p53 was confirmed by monitoring cell proliferation following treatment with an inhibitor that stabilizes p53 by preventing its ubiquitination by MDM2 (**Fig. S5A**). Analysis of stopwatch function partitioned the 15 p53-wildtype cell lines into 3 groups: 5 with a functional stopwatch, 2 with a partially compromised stopwatch, and 8 with an inactive stopwatch (**Fig. 4A,B; Fig. S5B-E**). In two neuroblastoma-derived p53-wildtype cancer cell lines, stopwatch activation led to significant cell death instead of arrest (**Fig. 4A; Fig. S5C**), presumably due to higher expression/activity of the cell death machinery. Of the 5 cell lines with a functional stopwatch, 3 appeared to have a higher temporal threshold for penetrant stopwatch activation (**Fig. 4A; Fig. S5C**).

Consistent with the prior bioinformatic analysis suggesting that *USP28* and *TP53BP1* exhibit a high frequency of deleterious mutations in cancer ((Davoli et al., 2013); **Fig. S4G,H**), mutations in *USP28* and/or *TP53BP1* are present in 4 of the 8 cell lines lacking stopwatch function (**Fig. 4C; Fig. S5F,G)**. Modeling mutations that resembled cancer-associated mutations in a nontransformed cell line revealed that, whereas frameshift mutations of both alleles of either *USP28* or *TP53BP1* eliminated protein expression and abrogated stopwatch function, heterozygous frameshift mutations reduced protein expression by ~50% and compromised stopwatch penetrance (**Fig. S6A-E)**. 3 other p53-wildtype cancer cell lines lacking stopwatch function had genetic alterations that dampen p53 signaling (**Fig. S6F-H**). In particular, mutations that truncate and hyperactivate the p53-antagonizing phosphatase WIP1 (Kleiblova et al., 2013) were associated with loss of stopwatch function, and treatment with a WIP1 inhibitor (Gilmartin et al., 2014) partially restored stopwatch activity (**Fig. S6F,G**). Collectively these results suggest that both p53-mutant and a significant proportion of p53-wildtype human cancers have compromised mitotic stopwatch function, which likely enables them to tolerate problematic mitoses that are both a cause and a consequence of cancer genomic instability.

### Stopwatch status predicts sensitivity to mitotic inhibitors

As the stopwatch monitors mitotic duration to control proliferation, its functional status has the potential to influence the efficacy of agents that prolong mitosis and are either currently used or in development as cancer therapeutic agents. To prolong mitosis to a moderate extent, we employed the PLK4 inhibitor centrinone (Wong et al., 2015) which delays spindle assembly due to loss of centrosomes (**Fig. S7A-D**). Monitoring the proliferation of the 3 p53-mutant and 15 p53-wildtype cancer cell lines revealed that p53-mutant lines continued to proliferate in PLK4i, albeit at reduced rates due to mitotic challenges cause by centrosome loss (**Fig. 4D**; **Fig. S7E** (Wong et al., 2015)). For the p53-wildtype cancer lines, proliferation in PLK4i was broadly inversely correlated with stopwatch status. p53-wildtype lines with a functional mitotic stopwatch exhibited rapid and penetrant cessation of proliferation in PLK4i (**Fig. 4D; Fig. S5H**); by contrast, those with a compromised or absent stopwatch exhibited improved proliferation in PLK4i (**Fig. 4D**). To extend this correlational analysis to an isogenic context, CHP134 neuroblastoma cells that possess an intact stopwatch and are highly sensitive to PLK4i were engineered to knockdown or mutate the 3 stopwatch complex subunits. Inactivation of the stopwatch significantly improved proliferation in PLK4i, relative to parental CHP134 cells (**Fig. 4D; Fig. S7F-H**), indicating that a functional stopwatch enhances the response to PLK4i treatment. Specific PLK4 inhibitors, while having the potential to target specific cancer types (Meitinger et al., 2020; Yeow et al., 2020), are currently tool compounds. Thus, we next assessed if stopwatch status impacted sensitivity to the widely used mitosis-targeting chemotherapy agent taxol and to a clinically-tested inhibitor of the mitotic kinesin CENPE (Bernabeu et al., 2017; Dumontet and Jordan, 2010; Penna et al., 2017; Qian et al., 2010; Tischer and Gergely, 2019). Low dose taxol and CENPEi treatments extend mitosis by mildly perturbing microtubule dynamics and chromosome congression, respectively. In both CHP134 neuroblastoma cells and in a nontransformed model cell line, mutation of stopwatch complex subunits significantly improved proliferation relative to the parental lines following low dose taxol or CENPEi treatment (**Fig. 4E, Fig. S7I**). Thus, stopwatch status influences the efficacy of therapeutic agents currently in use or being developed to target mitotic processes and could serve as a potential biomarker for their use in cancer treatment.

## Conclusion

Defects in chromosome segregation during mitosis, which not only generate cells with incorrect numbers of chromosomes but also precipitate drastic chromosome rearrangements, are intermediates during the generation of nearly all cancers (Garribba and Santaguida, 2022; Li and Zhu, 2022). Given the hundreds of billions of cells that divide in an adult human every day, monitoring these mitoses to filter out potentially problematic cells poses a significant challenge. The work described here elucidates how time spent in mitosis is monitored to control proliferation, via PLK1 kinase-dependent assembly of mitotic stopwatch complexes that are transmitted to daughter cells. The ability of this mechanism to detect subtle repeated extensions of mitosis functions as a fidelity filter in a proliferative cell population and explains its frequent inactivation in cancers, as well as the classification of core stopwatch complex components as tumor suppressors. Compromised stopwatch function is likely important for the tolerance of problematic mitoses that are contributors to and a consequence of the aneuploidy and genomic instability that is characteristic of cancers.

## Materials and Methods

### Chemical inhibitors

The chemical inhibitors used in this study were: centrinone (LCR-263; 150 nM; synthesized by Sundia MediTech; monastrol (100 μM; Tocris Bioscience); MDM2 inhibitor (Mdm2i; 1 μM; R7112; Proactive Molecular Research); BI2536 (PLK1i; 100 nM; MedChem Express); BI6727 (PLK1i; 25 nM; SelleckChem); GSK461364 (PLK1i; 25 nM; SelleckChem); MK-8745 (AuroraAi; 10 μM; SelleckChem); JQ1 (Bromodomain inhibitor; 10 μM; MedChem Express); CHIR124 (CHKi; 100 nM; Axon); CCT24533 (CHK2i; 1 μM; MedChem Express); CHK2 inhibitor II (CHK2i; 1 μM; Sigma-Aldrich); doxorubicin (Sigma-Aldrich); aphidicolin (1 μM; Cayman); BAY1895344 (ATRi; 1 μM; SelleckChem); AZD7648 (DNA-PKi; 1 μM; SelleckChem); AZD1390 (ATMi; 1 μM; Chemgood LLC); Palbociclib (CDK4/6i; 0.5 μM; SelleckChem); RO3306 (CDK1i; 10 μM; Calbiochem); GSK2830371 (WIP1i; 10μM; Seleckchem); paclitaxel (Taxol; Sigma-Aldrich); GSK923295 (CENPEi; SeleckChem).

### Antibodies

The Cep192 antibody (raised against aa 1-211; used at 0.5 μg/ml) was previously described (Wong et al., 2015). The following antibodies were purchased from commercial sources, with their working concentrations indicated in parentheses: anti-53BP1 (1:5,000; NB100-304; Novus Biologicals), USP28 (1:1,000; ab126604; Abcam), p53 (1:100; OP43; EMD Millipore), p21 (1:1,000; #2947; Cell Signaling Technology), anti-α-tubulin (1:5,000; DM1A; Sigma-Aldrich), anti-GAPDH (1:1,000; 14C10; Cell Signaling Technology), anti-Histone H3.3 (1:1,000; 09-838; EMD Millipore); anti-γH2AX (Ser139; 1:2000; 05-636; EMD Milipore). Secondary antibodies were purchased from Jackson ImmunoResearch and GE Healthcare.

### Cell lines

All cell lines used in this study are described in **Table S1**; the unique identifiers used for cell lines are described in *Fig. S1A*. RPE1 (hTERT RPE-1), U2OS, HCT116, CHP212, G401, SJSA1, RKO, U87, MCF7, BT549, A204 and A375 were obtained from the American Type Culture Collection (ATCC); CHP134 and DLD1 were obtained from Sigma-Aldrich (ECACC general collection) and CW2 line from Cell Bank Riken. LOX-IMVI was obtained from the NCI-60 collection. BT12, BT16 and SF8628 were a gift from Nalin Gupta (UCSF). Cell lines were cultured as recommended at 37°C and 5% CO_2_, supplementing the growth media with 100 IU/ml penicillin and 100 μg/ml streptomycin.

The RPE1 *USP28Δ, TP53BP1Δ, TP53sh* cell lines have been described (Meitinger et al., 2016; Meitinger et al., 2020). Knock-out and frameshift mutations were generated using transient transfection (RPE1 *USP28*Δ; RPE1 *TP53BP1*Δ; RPE1 *USP28^fs/fs^*; RPE1 *TP53BP1^fs/fs^*; RPE1 *CDKN1A*Δ; CHP134 *USP28^mut^;* CHP134 *TP53BP1*Δ) or nucleofection of preassembled RNP complexes (RPE1 *USP2^wts/fs^*; RPE1 *TP53BP1^wt/fs^*). Knockout and frameshift mutations were introduced using gene-specific sequences that target Cas9 to *USP28* (knockout targeting exon 4, TGAGCGTTTAGTTTCTGCAG; frameshift in exon 23, TGCTCTGGTAGGCATATACC), *TP53BP1* (knockout targeting exon 3, CTGCTCAATGACCTGACTGA; frameshift in exon 19, GTTTCCCCTTCACAGACTGG) or *CDKN1A* (knockout targeting exon 2, ATGTCCGTCAGAACCCATG). For transient transfections, double-stranded oligonucleotides for specific guide RNAs were cloned into PX459 [a gift from Feng Zhang (Addgene plasmid # 48139; http://n2t.net/addgene:48139; RRID:Addgene_48139)] (Ran et al., 2013). RPE1 cells were plated in 10 cm plates at 500,000 cells/plate the day before transfection. Cells were transfected with plasmid using Lipofectamine 3000 according to the manufacturer’s instructions (ThermoFisher). For nucleofection of RNP complexes 4 μl crRNA (100 μM) and 4 μl tracrRNA (100 μM) were incubated at 95°C for 5 min, cooled at room temperature for 15 min, incubated with 10 μl Cas9 (40 μM) for 20 min at room temperature, and nucleofected according to the manufacturer’s instructions (Lonza). For lentiviral infection, gRNAs were cloned into LentiCRISPR v2 (gift from Feng Zhang; Addgene plasmid # 52961; http://n2t.net/addgene:52961; RRID:Addgene_52961; (Sanjana et al., 2014)). Two days after transfection or nucleofection, 150 nM centrinone was added to select for mutants; control RPE1 cells and CHP134 cells arrest and die, respectively, after 3-4 divisions in centrinone. After 10 days, centrinone-resistant RPE1 or CHP134 cells were plated at low density in 96 well plates (30-40 cells per plate) for single clone selection. For selecting CHP134 *TP53BP1*Δ and USP28^mut^ mutants, centrinone-resistant CHP134 cells were plated at a density that supported direct picking of clones from 10 cm plates. Gene knock-out was assessed by genotyping of the sequence surrounding the CRISPR cut site (Brinkman et al., 2014) and/or by immunoblotting.

RPE1 *p21-mNeonGreen* cells were generated using CRISPR/Cas9 in combination with rAAV mediated delivery of the repair construct as previously described (Kaulich and Dowdy, 2015). The gRNA, designed to cut close to the stop codon of CDKN1A(*p21*) (TTTGAGGCCCTCGCGCTTCC), was cloned into PX459 ((Ran et al., 2013); gift from Feng Zhang; Addgene plasmid # 48139; http://n2t.net/addgene:48139; RRID:Addgene_48139). The repair construct, cloned into pSEPT, contained NeonGreen for C-terminal fusion to p21 linked through a P2A sequence to the neomycin resistance gene Tn5 aminoglycoside phosphotransferase; this cassette was flanked on either side by homology arms corresponding to the regions up and down stream of the gRNA target site. The left homology arm contained 926 bp upstream of the stop codon and the right homology arm contained 684 bp downstream of the gRNA including 6bp overlapping with the gRNA sequence.

The following transgenes were stably integrated into the genome using lentiviral constructs (**see Table S2**): H2B-mRFP (EF1alpha promoter); TRE3pro-Cas9 (Edit-R Inducible Lentiviral Cas9; Dharmacon); 3FLAG-APOBEC-1-Cas9(D10A)-UGi (Base Editor;(Zafra et al., 2018)). Cell lines for inducible knock-out of Comet or induction of a double strand break at the control *AAVS1* locus were generated as previously described (Meitinger et al., 2020). In brief, cell lines were generated by sequential lentiviral integration of inducible Cas9 and a Comet sgRNA expressing plasmid based on the lentiGuide-Puro plasmid ((Sanjana et al., 2014); gift from Feng Zhang; Addgene plasmid # 52963; http://n2t.net/addgene:52963; RRID: Addgene_52963). The Comet gRNA (AAGTGCTTAAGCTGTTCATA) targets exon 3 and the *AAVS1* gRNA (GGGCCACTAGGGACAGGAT) targets the intron between exon 1 and 2 of *PPP1R12C*. Cas9 expression was induced with 1 μg/ml doxycycline.

For base editing, Cas9 from the lentiCas9-Blast plasmid ((Sanjana et al., 2014); gift from Feng Zhang; Addgene plasmid # 52962; http://n2t.net/addgene:52962; RRID:Addgene_52962) was replaced with 3FLAG-APOBEC-1-Cas9(D10A)-UGI from the pLenti-FNLS-P2A-Puro plasmid ((Zafra et al., 2018); gift from Lukas Dow; Addgene plasmid # 110841; http://n2t.net/addgene:110841; RRID:Addgene_110841). After lentiviral integration of 3FLAG-APOBEC-1-Cas9(D10A)-UGI, a specific gRNA (TATGTCCTTTCACCACTCCT) designed to facilitate mutation of 53BP1 Gly1560 to Lys was integrated using the lentiGuide-Puro plasmid (Sanjana et al., 2014)).

Viral particles were generated by transfecting the lentiviral construct into HEK-293T cells using Lenti-X Packaging Single Shots (Clontech). 48 hours after transfection, virus-containing culture supernatant was harvested and added to the growth medium of cells in combination with 2.5-8 μg/ml polybrene (EMD Millipore). Populations of each cell line were selected by FACS or antibiotics (Blasticidin, 5 μg/ml; Neomycin, 400 μg/ml; Puromycin, 10 μg/ml for RPE1, 0.5 μg/ml for CHP134). Single clones were isolated in 96-well plates.

The RPE1 cell line expressing an S-phase marker was generated using the lentiviral CSII-EF mVenus-hGeminin(1-110) construct (Sakaue-Sawano et al., 2011). Virus production and cell transfection was done as described above. mVenus-hGeminin (1-110) expressing cells were sorted by FACS. The BI6727/BI2536-resistant mutation R136G in PLK1 was generated by chemical transfection of a specific gRNA (TGTTGGAGCTCTGCCGCCGG) expressing plasmid (pX459) and a repair oligonucleotide (GCGCTCCAACTGCCCCGCAGGCAGTTCCAGTTCCCCAGCAGCGACACTCACCCCGCGGCGGCAGAGCTCCAACACCACGAACACGAAGTCGTTGTCCTCG). The gRNA targets a region close to the mutation site. pSpCas9(BB)-2A-Puro (PX459) V2.0 was a gift from Feng Zhang ((Ran et al., 2013); Addgene plasmid # 62988; http://n2t.net/addgene:62988; RRID:Addgene_62988). Positive clones were selected with 25 nM BI6727 two days after the chemical transfection.

### Immunofluorescence

For immunofluorescence, ~10,000 cells per well were seeded into 96-well plates one day before fixation. Centrosome depletion was monitored after 8 days of centrinone treatment with the CEP192 antibody. Cells were fixed in 100 μl ice-cold methanol for 7 minutes at −20°C. Cells were washed twice with washing buffer (PBS containing 0.1% Triton X-100) and blocked with blocking buffer (PBS containing 2% BSA, 0.1% Triton X-100 and 0.1% sodium azide) overnight. After blocking, cells were incubated for 1-2 hours with primary antibody in fresh blocking buffer (concentrations as indicated above). Cells were washed three times with washing buffer, prior to 1-hour incubation with the secondary antibody and DNA staining Hoechst 33342 dye. Finally, cells were washed three times with washing buffer. Images were acquired on a CV7000 spinning disk confocal system (Yokogawa Electric Corporation) equipped with a 40X (0.95 NA) or a 60X (water, 1.2 NA) U-PlanApo objective and a 2560×2160 pixel sCMOS camera (Andor). Image acquisition was performed using CV7000 software.

### Live cell imaging

Live cell imaging was performed on the CQ1 spinning disk confocal system (Yokogawa Electric Corporation) equipped with a 40X 0.95 NA U-PlanApo objective and a 2560×2160 pixel sCMOS camera (Andor) at 37°C and 5% CO_2_. Image acquisition and data analysis were performed using CQ1 software and ImageJ, respectively.

#### Mitotic stopwatch assay

All imaged cell lines were engineered to express H2B-RFP (see **Table S1**). One day before imaging, cells were seeded into 96-well cycloolefin plates at 2,000-4,000 cells/well. On the day of the experiment, asynchronous cells were first imaged for 4-6h in 100 μM monastrol. During monastrol treatment, cells enter at different times into mitosis and delay in prometaphase. This generates a set of mother cells that experience different mitotic durations prior to monastrol washout. Images of H2B-RFP were acquired with 5 x 2 μm z-sections in the RFP channel (25% power, 150 ms) at 10-minute intervals. After monastrol washout, the mother cells complete mitosis and divide. The resulting daughter cells were imaged at 10-minute intervals and tracked for up to 72h. The fate of the daughter cells was classified into ‘arrest’, ‘death’ or ‘proliferate’. Inhibitors were added during prolonged mitosis and washed out together with monastrol, unless noted otherwise.

#### Measurement of Mitotic duration

Mitotic duration was measured in cell lines expressing H2B-RFP (see **Table S1**). To induce prolonged mitosis cells were either treated with doxycycline to induce Comet knockout (see generation of inducible knockout mutants above) or centrinone to deplete centrosomes (*Fig. 1E,F; Fig. S2A,B*). Doxycycline and centrinone were administered 1-4 days before imaging as indicated for each experiment. To determine the consequences of centrosome depletion on the length of mitosis in different cancer cell lines (*Fig. S7D*), cells were treated for 3 cell cycles with 150 nM centrinone to deplete centrosomes or with DMSO as a control. Mitotic duration was quantified as the time from nuclear envelope breakdown (NEBD) until chromosome decondensation. Prior to this analysis, the cell cycle duration for all analyzed cancer cell lines was measured by H2B-RFP live imaging of each cell line and quantifying time from NEBD of a mother cell to NEBD of its daughters. For all experiments, cells were seeded one day before imaging into 96-well cycloolefin plates at 5,000-10,000 cells/well. Images of H2B-RFP were acquired with 5 x 2 μm z-sections in the RFP channel (25% power, 150 ms) at 4-5-minute intervals.

#### Monitoring p21 expression

To measure p53 activation after prolonged mitosis, the coding sequence for mNeonGreen was fused to the p53 target and effector p21 (encoded by *CDKN1A*) at both endogenous *CDKN1A* loci in RPE1 cells. Mitosis was prolonged using temporal monastrol or centrinone treatment as described above. H2B-RFP was used to track cells and mark the nuclear area throughout the experiment. Cells were seeded one day before imaging into 96-well cycloolefin plates at 4,000 cells/well. Images of p21-mNeonGreen (GFP channel, 35% power, 150 ms) and H2B-RFP (RFP channel, 25% power, 150 ms) were acquired with 5 x 2 μm z-sections at 10-minute intervals during monastrol treatment and at 20-minute intervals after monastrol washout or during centrinone treatment. Expression of p21 was quantified in the nucleus at each timepoint using Fiji software (Schindelin et al., 2012). The normalized signal intensity was determined by drawing a rectangle around the nucleus (w x h) and quantifying the average p21-mNG signal. The average background was determined by drawing a rectangle 2 pixels larger in width and height and quantifying the average p21-mNG signal in the intervening region. The total intensity was the average intensity in the smaller rectangle minus the background intensity multiplied by the area of the smaller rectangle.

To measure G1 duration after prolonged mitosis, cells were engineered to express the cell cycle marker mVenus-hGeminin and the nuclear marker H2B-RFP. Mitosis was prolonged using monastrol treatment as described above. H2B-RFP was used to track cells and mark the nuclear area throughout the experiment. Cells were seeded one day before imaging into 96-well cycloolefin plates at 4,000 cells/well. Images of mVenus-hGeminin (GFP channel, 35% power, 150 ms) and H2B-RFP (RFP channel, 25% power, 150 ms) were acquired with 5 x 2 μm z-sections at 10-minute intervals during monastrol treatment and at 20-minute intervals after monastrol washout. G1 duration was determined as the time between chromosome decondensation and the start of mVenus-hGeminin expression (S-phase).

### Proliferation and viability assays

For proliferation analysis, cells were seeded into 6 well plates in triplicate at 25,000-150,000 cells/well and treated with the indicated inhibitors or DMSO as a control. At 72-hour or 96-hour intervals, cells were harvested, counted and, for passaging assays, re-plated at 25,000-150,000 cells/well. Cell counting was performed using a TC20 automated cell counter (Bio-Rad).

### DNA damage analysis

ATM and ATR inhibitors prevent the formation of γH2AX foci at DNA damage sites, whereas DNA-PK inhibition at low concentrations of doxorubicin leads to an accumulation of γH2AX foci (Durant et al., 2018; Fok et al., 2019; Kurose et al., 2006). To assess inhibitor efficacy, 4-8,000 cells per well were seeded into 96-well plates one day prior treatment with either doxorubicin or aphidicolin, at the indicated concentrations, along with ATM and ATR inhibitors for 2 hours (for ATMi & ATRi analysis), or with DNA-PKi for 24 hours. Cells were prepared for immunostaining as described above and stained with a γH2AX-antibody to visualize DNA damage sites and Hoechst 33342 for DNA staining. Images were acquired on a CV7000 spinning disk confocal system (Yokogawa Electric Corporation) equipped with a 40x (0.95) U-PlanApo objective and a 2,560 x 2160-pixel sCMOS camera (Andor Technology). Quantification was performed using CV7000 analysis software to count the number of γH2AX foci within the nucleus for each cell. Each condition was analyzed in triplicate.

### Immunoblotting

For immunoblotting, cells were cultured in 10 cm plates, harvested at 50-80% confluence and lysed by sonication in RIPA buffer (Cell Signaling Technology; 20 mM Tris-HCl (pH 7.5), 150 mM NaCl, 1 mM Na_2_EDTA, 1 mM EGTA, 1% NP-40, 1% sodium deoxycholate, 2.5 mM sodium pyrophosphate, 1 mM beta-glycerophosphate, 1 mM Na_3_VO_4_, 1 μg/ml leupeptin) plus protease and phosphatase inhibitor cocktail (Thermo Fisher Scientific). Cell extracts were stored at −80°C until use. Extract concentrations were normalized based on protein concentration (Bio-Rad Protein Assay) and 20-30 μg protein was loaded per lane on Mini-PROTEAN gels (Bio-Rad) and transferred to PVDF membranes using a TransBlot Turbo system (Bio-Rad). Blocking and antibody incubations were performed in TBS-T + 5% non-fat dry milk. Detection was performed using HRP-conjugated secondary antibodies (GE Healthcare) with SuperSignal West Femto (Thermo Fisher Scientific) substrates. Membranes were imaged on a ChemiDoc MP system (Bio-Rad).

### Immunoprecipitation and cell lysate fractionation

For analysis of cells at particular cell cycle stages, cells were arrested in G1 by treatment with 0.5 μM Palbociclib, in G2 by treatment with 10 μM RO-3306, and in Mitosis by treatment with 200 ng/ml Nocodazole or 100 nM BI2536 for 14-20 hours. DNA damage was induced using 1μM doxorubicin treatment for 16 hours.

For immunoprecipitation assays, ~20 million cells were harvested and washed with PBS. Cells were resuspended in lysis buffer (20 mM Tris/HCl pH 7.5, 50-200 mM NaCl, 0.5% Triton X-100, 5 mM EGTA, 1 mM dithiothreitol, 2 mM MgCl_2_ and EDTA-free protease inhibitor cocktail (Roche)) and lysed in an ice-cold sonicating water bath for 5 minutes. All immunoprecipitation assays were performed using 50 mM NaCl unless otherwise noted. After 15-minute centrifugation at 15,000 x g and 4 °C, soluble lysates were collected, and protein concentrations were quantified. Equal amounts of lysates were incubated with anti-53BP1 for 2 hours at 4°C and subsequently with Protein A magnetic beads (Thermo Fisher Scientific) for 1 hour at 4 °C. The beads were washed five times with lysis buffer and resuspended in SDS sample buffer. For immunoblotting, equal volumes of samples were run on Mini-PROTEAN gels (Bio-Rad) and transferred to PVDF membranes using a TransBlot Turbo system (Bio-Rad). Blocking and antibody incubations were performed in TBS-T plus 5% nonfat dry milk. Immunoblotting was performed as described above. For immunoprecipitation assays following DNA damage, cells were treated for 16 hours with 1 μM doxorubicin. The lysis buffer contained 200 mM NaCl to increase the solubility of 53BP1. As control for 53BP1 and USP28 interaction, mitotic cells were harvested as described above and treated with the same lysis buffer containing 200 mM NaCl. Immunoprecipitation assays were performed as described above.

For immunoprecipitation assays in G1 phase following prolonged mitosis, cells were first arrested in mitosis with 200 ng/ml Nocodazole with and without 100 nM of the PLK1 inhibitor BI2536 for 14h. To allow cells to exit mitosis and enter G1 phase, mitotic cells were directly washed of the plate, washed three times with PBS in a conical flask and replated onto 15 cm dishes. Plated cells were allowed to exit mitosis and enter G1 phase. After 6 hours G1 phase cells were harvested, and immunoprecipitation assays were performed as described above.

For cell lysate fractionation, cells were cultured on 15 cm dishes. To induce DNA damage, cells were treated for 16h with doxorubicin (1 μM), and to arrest cells in mitosis cells were treated for 16 with Nocodazole (0.83 μM). Four plates were harvested for each condition. Control and doxorubicin treated cells were harvested with trypsin and washed with PBS. Mitotic cells were directly washed of the plates. All cells were washed once with PBS and resuspended in 5 ml PBS. Suspended cells were counted. Cells were divided into aliquots of 1.5 million cells, pelleted and resuspended with 200 μl lysis buffer containing different amounts of sodium chloride (50 mM, 100 mM, 150 mM and 200 mM). Cells were sonicated for 5 min to generate the whole cell extract. 100 μl whole cell extract of each sample was collected for immunoblotting. The remaining 100 μl WCE was centrifuged for 15 minutes at 15,000 x g at 4 °C to generate the supernatant and pellet fractions. Pellets were resuspended in lysis buffer to equal the total volume of the supernatant. 40μl 4x sample buffer was added to each sample. Equal amounts of whole cell extract, supernatant and pellet fractions were analyzed by immunoblotting.

## Acknowledgments

We thank Midori Ohta and Aleesa Schlientz for feedback on the manuscript and members of the Oegema and Desai labs for helpful discussion.

## Author contributions

Conceptualization, FM, AS, AD, KO

Methodology, FM, RD, MM

Resources, AS, AD, KO

Investigation, FM, RD, AS, KO, AD

Writing – Original Draft, FM, KO, AD

Writing – Review & Editing, FM, AS, RD, KO, AD

Funding, KO, AD

## Data and materials availability

All data and materials are available in the main text or the supplementary materials or upon request.

**Fig. 1.**
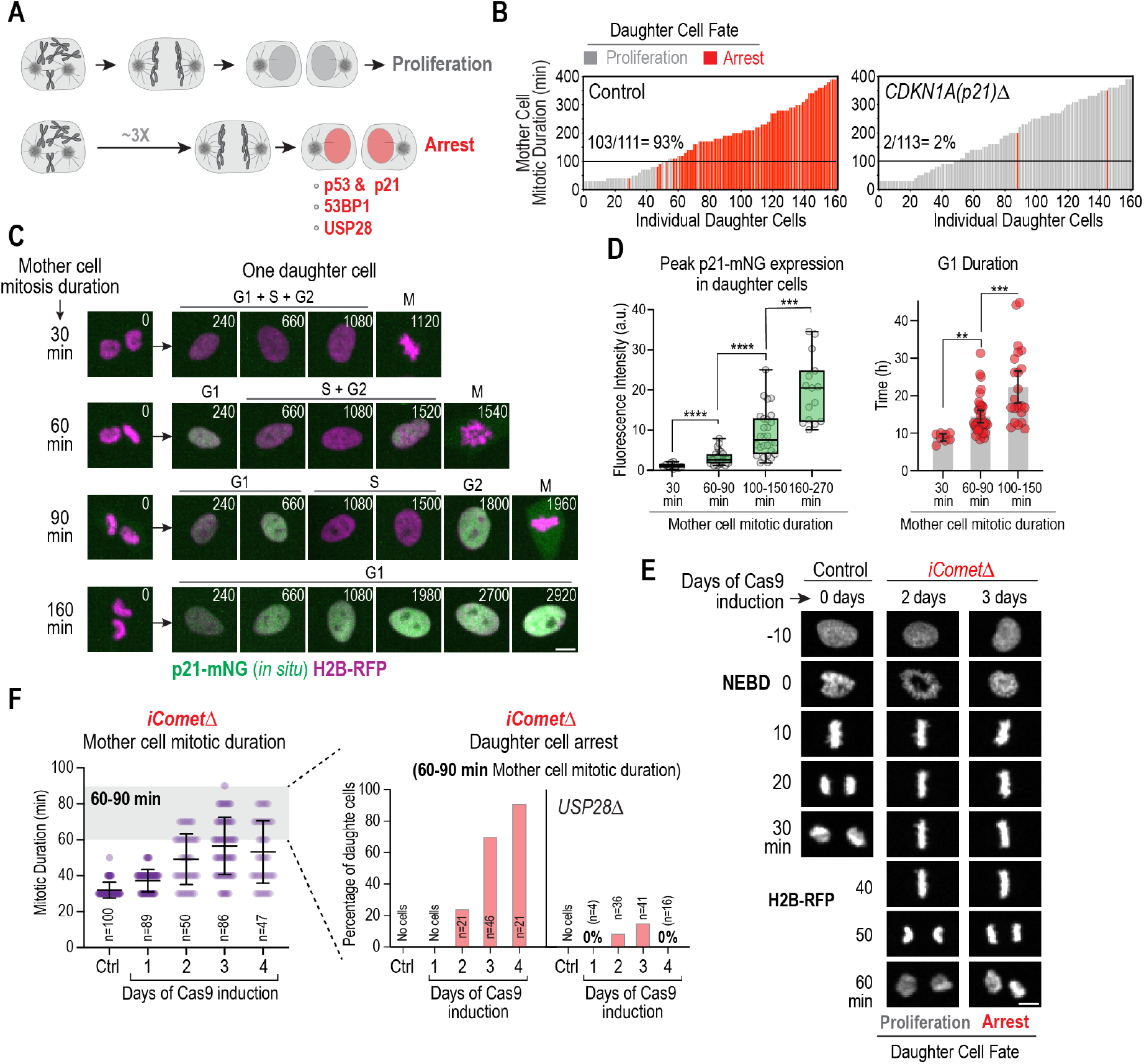
Control of G1 progression and cell proliferation by memories of mitosis. (**A**) Schematic of the mitotic stopwatch. Prolonging mitosis beyond a threshold time leads to a G1 arrest of daughter cells that requires p53, p21, 53BP1 and USP28. (**B**) Plots showing the results of mitotic stopwatch assays. Transient disruption of spindle assembly generates mother cells of varying mitotic durations whose daughters’ fates are tracked by live imaging. Each bar represents a daughter cell, with bar height representing the mitotic duration of its mother and color representing its fate (proliferation or arrest). Percentage of daughters of mothers that spent ≥100 minutes in mitosis that arrested is noted above the black lines. See also *Fig. S1A,B*. (**C**) Stills from timelapse movies monitoring *in situ*-tagged p21-mNeonGreen and H2B-RFP. Panels show mother cells of different mitotic durations (*left*) and one of their daughters (*right*). (**D**) (*left*) Box-and-whiskers plot of peak daughter cell p21-mNG expression in G1, as a function of mother cell mitotic duration. (*right*) G1 duration of daughters produced by mother cells of the indicated mitotic durations. Mean and 95% CI are indicated. p-values in (C) and (D) are from t-tests (**:p<0.01; ***: p<0.001; ****: p<0.0001). See also *Fig. S1C-E*. (**E**) Panels from time-lapse movies of representative control and *iComet*Δ cells expressing H2B-RFP. NEBD: nuclear envelope breakdown. (**F**) (*left*) Mitotic duration at different days after induction of Comet knockout; Mean and SD are indicated. (*right*)Frequency of daughter cells that arrest produced by mothers with 60-90 min mitotic duration on different days following induction of Comet knockout. The Comet knockout was also induced and analyzed in *USP28*Δ and *TP53sh* cells; *USP28*Δ is shown. See also *Fig. S1F-I*. Scale bars in (C) & (E), 5 μm.

**Fig. 2.**
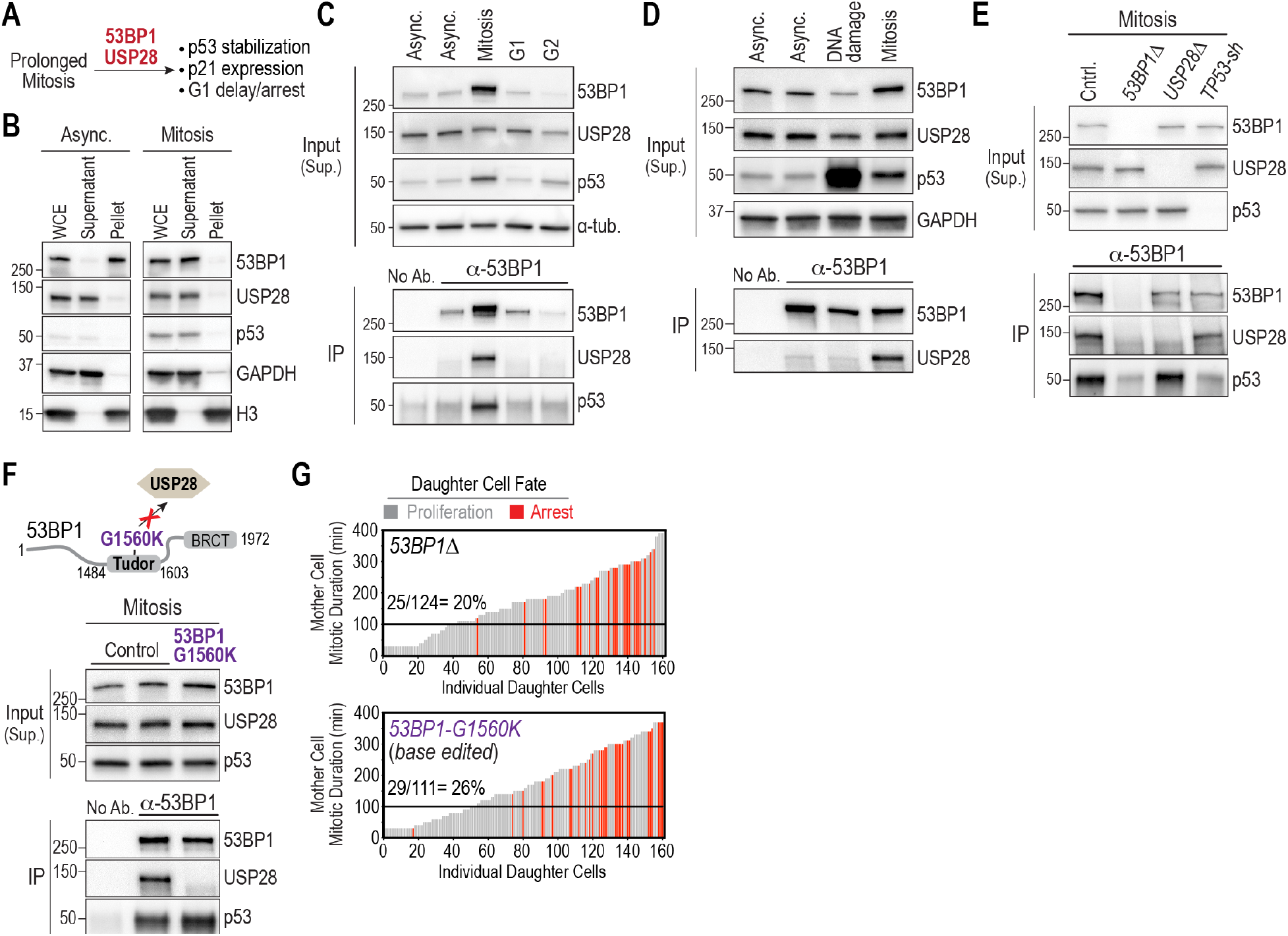
Mitosis-specific formation of stopwatch complexes transmits memory of mitotic duration to daughter cells. (**A**) Schematic highlighting the requirement for 53BP1 and USP28 in the mitotic stopwatch. (**B**) Immunoblots monitoring solubility in asynchronous and mitotic cell extracts. WCE: Whole Cell Extract. GAPDH and histone H3 are soluble and chromatin-bound insoluble controls, respectively. (**C**)-(**E**) Immunoblots of 53BP1 immunoprecipitates for the indicated conditions; a-tubulin and GAPDH serve as loading controls in (C) and (D), respectively. See also *Fig. S3A,B*. (**F**) (*top*) Schematic showing the location of a point mutation in 53BP1’s Tudor domain that disrupts interaction with USP28 (22). (*bottom*) 53BP1 immunoprecipitation analysis for the indicated conditions. (**G**) Mitotic stopwatch assays of engineered mutant cell lines. See also *Fig. S3C*.

**Fig. 3.**
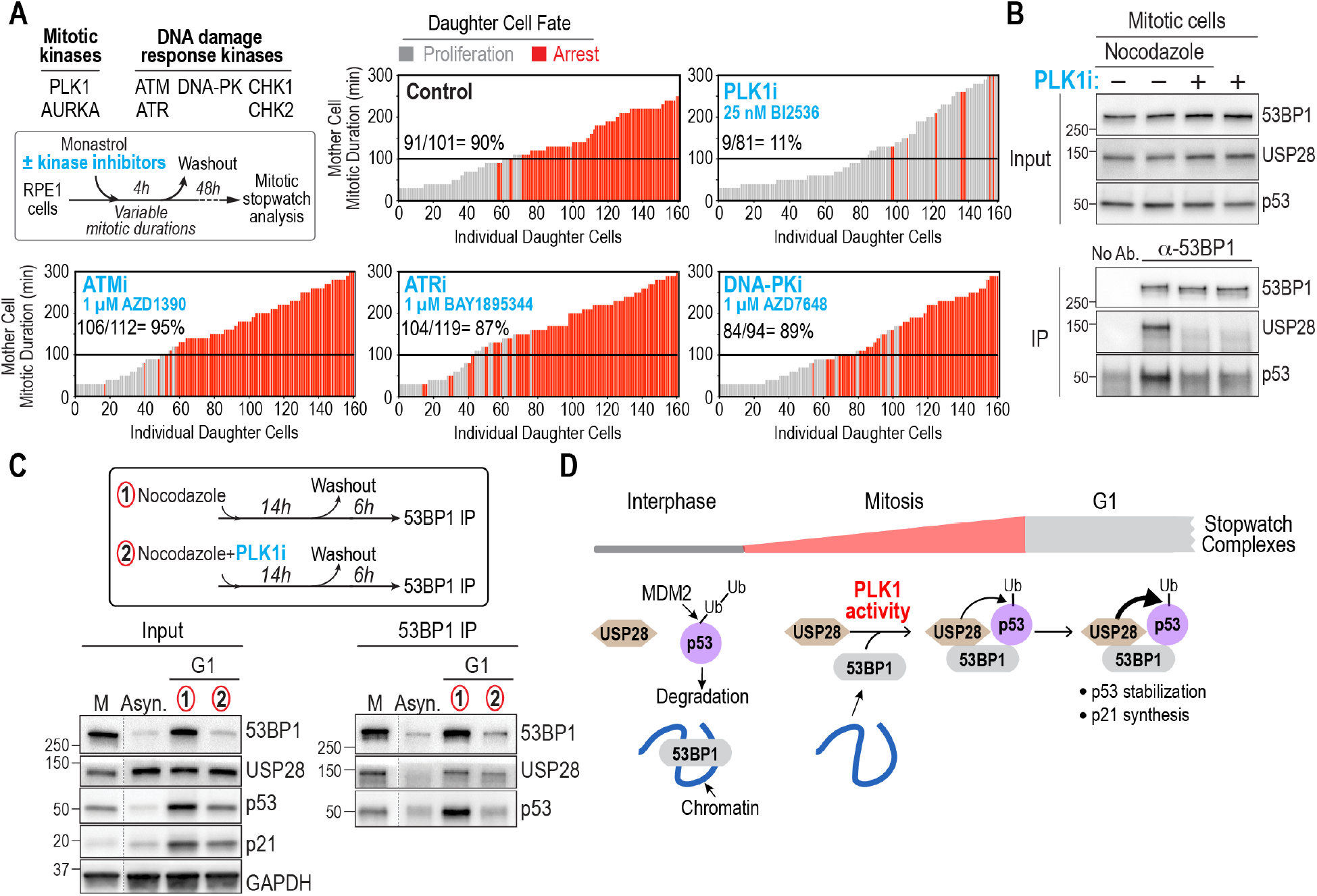
PLK1 kinase activity is central to the formation of stopwatch complexes that are transmitted to daughter cells. (**A**) (*top left*) Experimental approach and list of inhibited mitotic and DNA damage kinases. Plots show results of mitotic stopwatch assays for the indicated conditions. See also *Fig. S4A-F*. (**B**) 53BP1 immunoprecipitation analysis for the indicated conditions. (**C**) (*top*) Schematic of analysis examining the persistence of stopwatch complexes into G1. (*bottom*) Immunoblots of inputs and 53BP1 immunoprecipitates. M: Mitosis; Asyn.: asynchronous. GAPDH serves as a loading control. (**D**) Schematic showing the PLK1-dependent formation during mitosis of stopwatch complexes that persist into G1 where, depending on their abundance, they either trigger proliferation arrest or impart memory of prolonged mitosis into the next cell cycle.

**Fig. 4.**
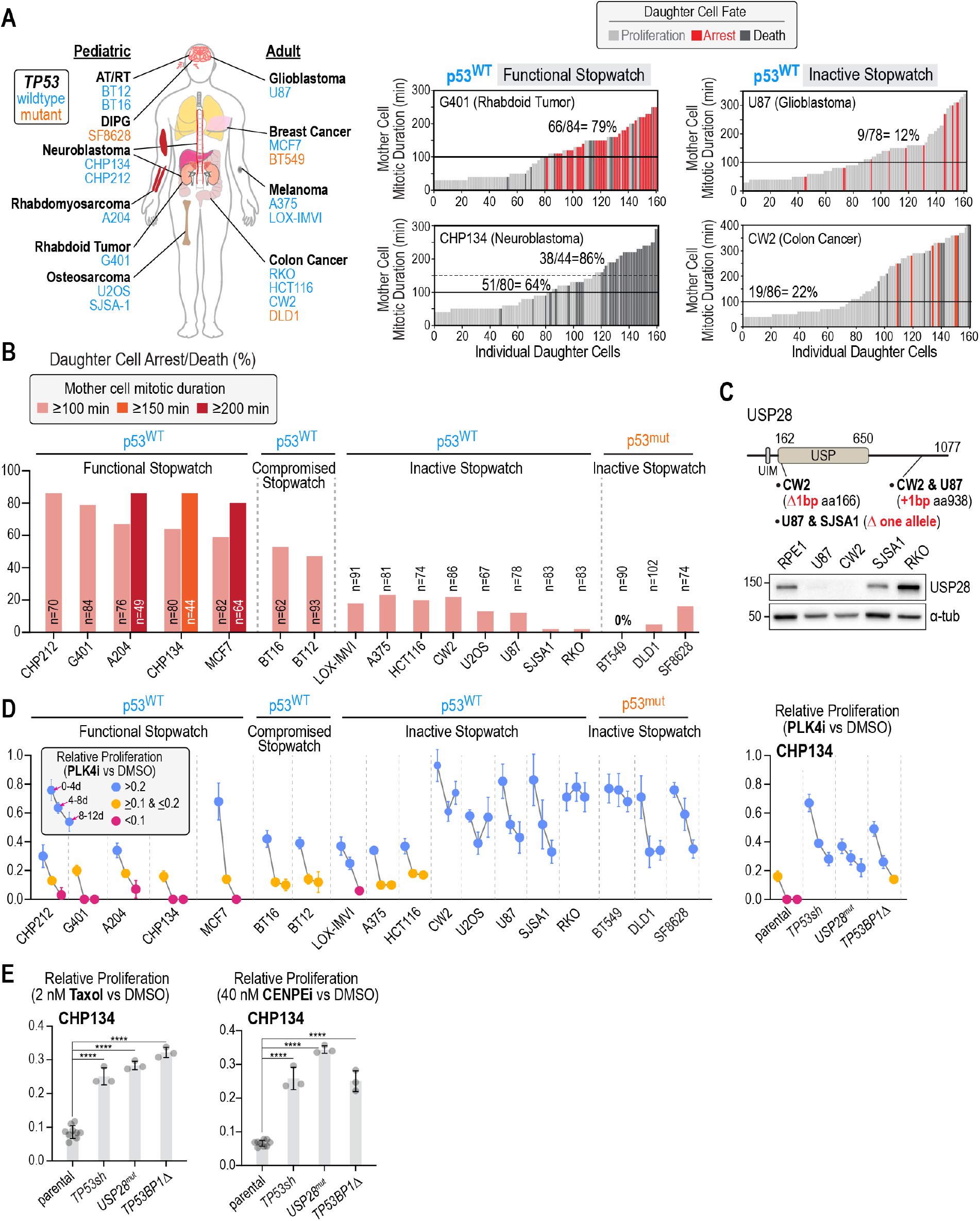
The mitotic stopwatch is compromised in cancers and influences efficacy of antimitotic agents. (**A**) (*left*) Schematic of diverse tissue-of-origin cancer-derived cell lines annotated as expressing wildtype (*blue*) or mutant (*orange*) p53; for experimental confirmation of p53 status see *Fig. S5A*. (*right*) Representative plots for p53-wildtype cancer-derived cell lines with functional or inactive mitotic stopwatches. See also *Fig. S5B-E*. (**B**) Plot quantifying percent daughter cell arrest/death from mitotic stopwatch plots for mother cell mitotic durations above the indicated thresholds in the p53-wildtype and p53-mutant cancer cell lines. (C) (*top*) USP28 schematic showing mutations identified in p53-wildtype cancer lines that lack stopwatch function. (*bottom*) USP28 immunoblot for the indicated cell lines; a-tubulin is a loading control. See also *Fig. S5F,G*. (**D**) (*left*) Mean relative proliferation in PLK4i versus DMSO is plotted for successive 4-day intervals in a passaging assay for 15 p53-wildtype and 3 p53-mutant cancer lines; see also *Fig. S5H*. p53-wildtype lines are grouped based on their stopwatch status. (*right*) Mean relative proliferation in PLK4i of CHP134 neuroblastoma cells and derived isogenic clonal lines with mutations/knockdown of stopwatch components. Error bars are the SD (n=3). (**E**) Mean relative proliferation of parental CHP134 cells and derived lines in 2 nM Taxol (*left*) or 40 nM CENPEi (*right*); n=9 for controls and 3 for other conditions. Error bars are the SD.

**Fig. S1.**
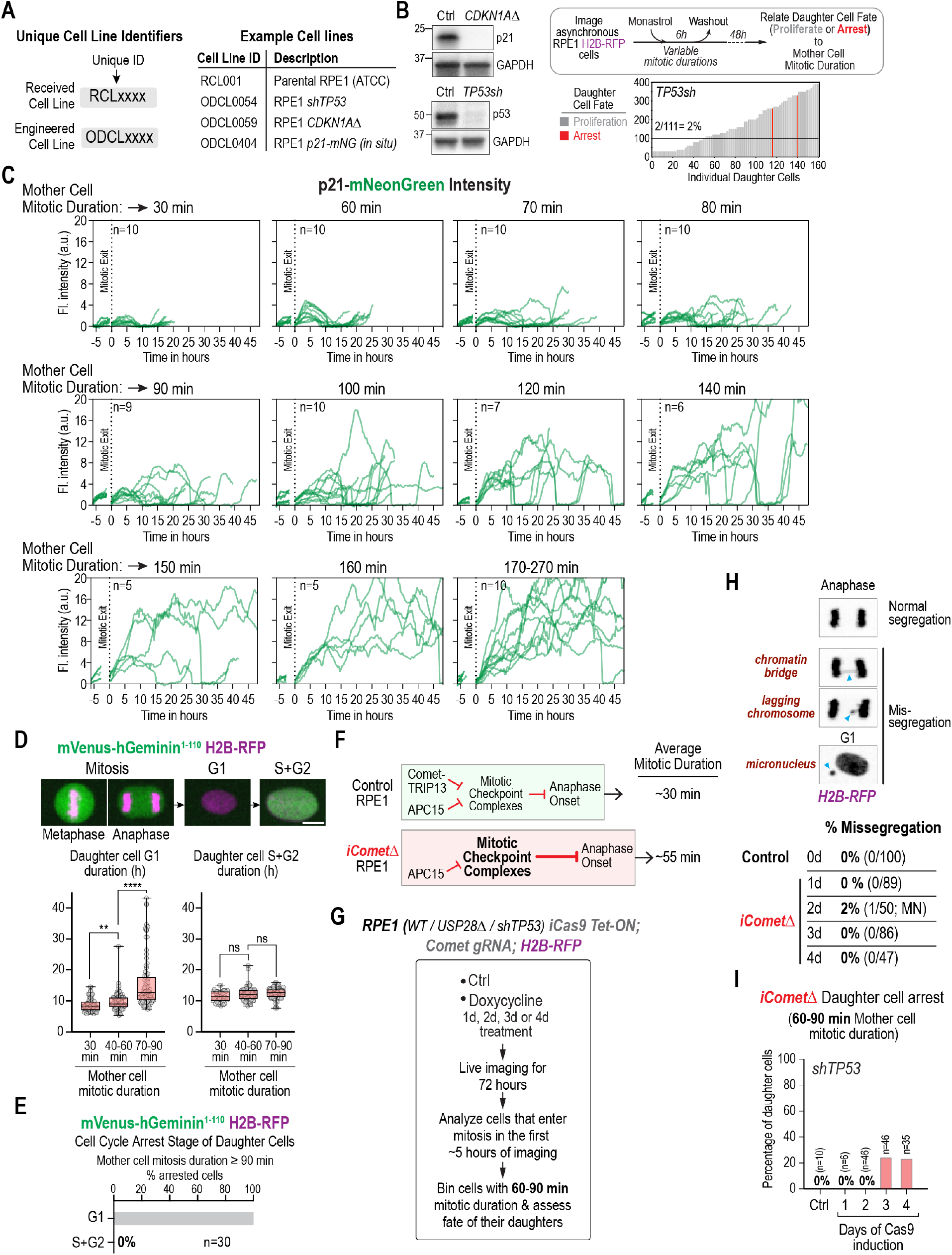
Cell line identifiers and immunoblots, mitotic stopwatch assay, p21-mNG expression traces, analysis of cell cycle phases, and Comet-KO details. (**A**) Depiction of unique cell line identifiers used in *Table S1* with specific examples. (**B**) (*left*) Immunoblots of control and indicated engineered RPE1 cell lines. GAPDH serves as a loading control. (*top right*) Schematic of mitotic stopwatch assay. Transient disruption of spindle assembly in an asynchronous culture, using the reversible mitotic kinesin inhibitor monastrol, generates mother cells with variable mitotic durations that produce daughters whose fate is tracked by live imaging. (*bottom right*) Mitotic stopwatch assay plot for p53-knockdown RPE1 cells; control RPE1 plot is in *Fig. 1B*. (**C**) Traces of nuclear p21-mNG fluorescence intensity in daughter cells produced by mother cells with the indicated mitotic durations. Dashed vertical line marks mitotic exit (t=0). (**D**) Box-and-whiskers plot of G1 and S+G2 durations as a function of mother cell mitotic duration, measured in an RPE1 cell line expressing a green-fluorescent geminin fragment and red-fluorescent H2B. Scale bar, 5 μm. p-values are from t-tests (ns: not significant; **:p<0.01; ***: p<0.001; ****; p<0.0001). (**E**) Cell cycle arrest stage after extended mitosis (≥90 min) in the RPE1 cell line expressing the green-fluorescent geminin fragment and red fluorescent histone. All daughter cells arrested in G1. (**F**) Schematic illustrating the two parallel mechanisms that disassemble mitotic checkpoint complexes that delay anaphase onset. Chromosomes not attached to spindle microtubules prolong mitosis by catalyzing formation of mitotic checkpoint complexes that inhibit the anaphase promoting complex/cyclosome (APC/C), the E3 ubiquitin ligase that triggers anaphase onset (51). Comet (also known as p31) accelerates APC/C activation and mitotic exit by targeting mitotic checkpoint complexes for disassembly by the AAA+ enzyme TRIP13 (16). Comet-TRIP13 is one of two mechanisms inactivating mitotic checkpoint complexes (17); thus, Comet inhibition prolongs metaphase but does not prevent mitotic exit. (**G**) Schematic of experiment in Fig. 1F. A gRNA targeting MAD2L1BP (which encodes Comet) was introduced into RPE1 cell lines (wildtype, USP28Δ or TP53sh) with doxycycline-inducible Cas9 that also express RFP-H2B. Comet knockout was induced by doxycycline addition for 1, 2, 3 or 4 days, after which cells were imaged for 72h. Cells that entered mitosis in the first 5h of filming were analyzed to measure their mitotic duration and assess daughter cell fate. (**H**) (*top*) Examples of normal or different types of defective chromosome segregation (blue arrowheads) in anaphase or early G1 RPE1 cells expressing H2B-RFP. (*bottom*) Frequency of missegregation events in iCometΔ RPE1 cells. No significant elevation of missegregation was observed following induction of Comet knockout. (**I**) Analysis of inducible Comet knockout in RPE1 TP53sh cells. Plot of the fate of daughters produced by mother cells with 60-90 min mitotic duration; results for the wildtype and USP28Δ RPE1 cell lines are in *Fig. 1F*.

**Fig. S2.**
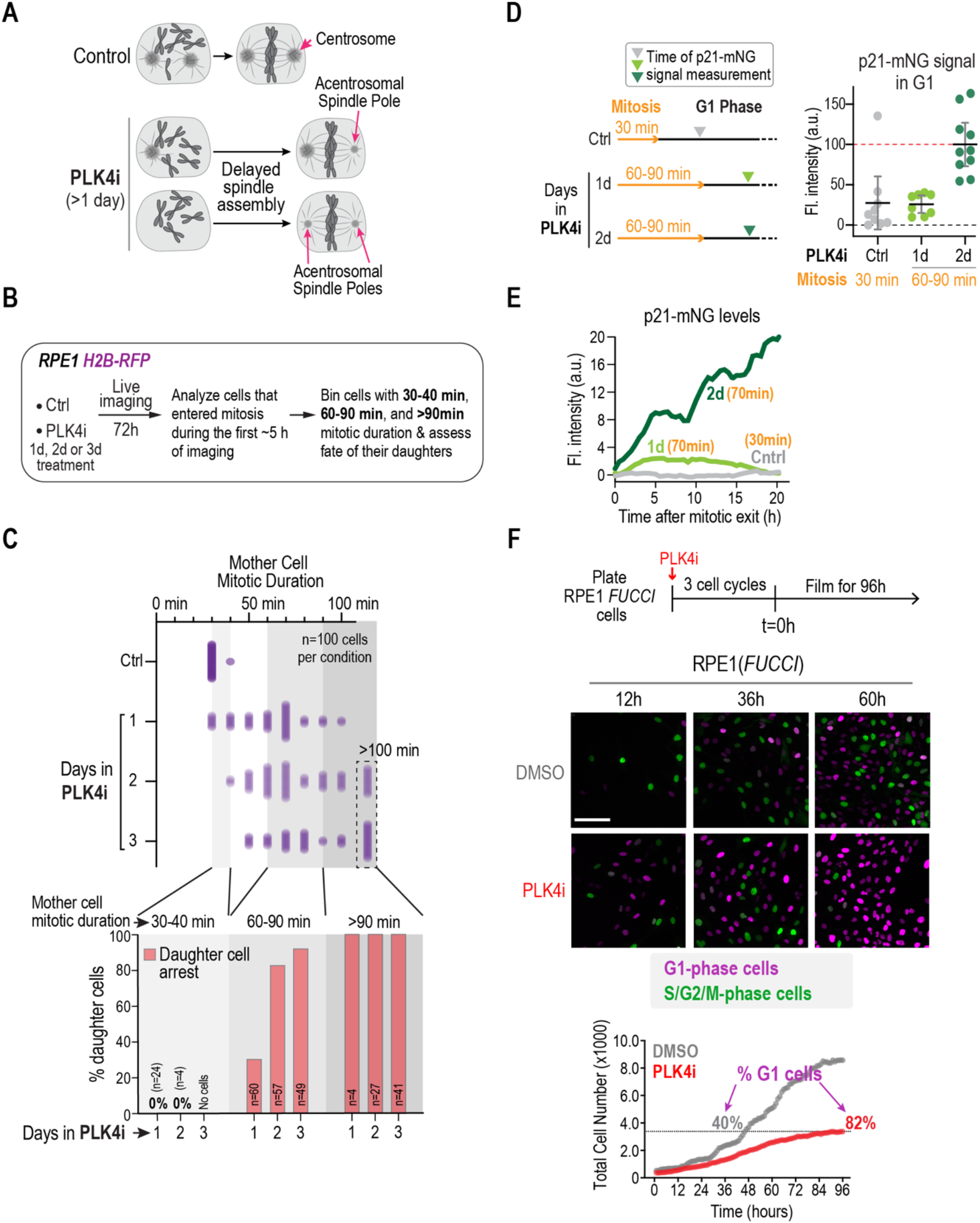
Evidence for multi-generational memory in the mitotic stopwatch from PLK4i treatment-induced extended mitosis. (**A**) Schematic illustrating the effect of centrosome loss, induced by treatment with the PLK4 inhibitor (PLK4i) centrinone (31), on mitotic duration. Loss of one or both centrosomes slows spindle assembly. (**B**) Schematic of the experiment used to monitor the effect of PLK4i over multiple cell cycles. RPE1 H2B-RFP cells were treated for 1d with DMSO vehicle or with PLK4i for 1, 2 or 3 days, and then imaged live for 72h. Cells that entered mitosis in the first 5h of filming were analyzed to measure their mitotic duration and to determine the fate of their daughters. Daughter cell fate was plotted relative to the indicated binned mitotic durations of their mothers. (**C**) (*top*) Graph plotting the distribution of mother cell mitotic durations for cells treated with PLK4i for the indicated number of days. Control is RPE1 cells treated for 1d with DMSO vehicle. (*bottom*) For the indicated ranges of mother cell mitotic duration (30-40 minutes; 60-90 minutes; >90 minutes), graph plots the percentage of daughter cells that arrested in G1 as a function of days of exposure to PLK4i. Daughters of cells that spent >90 minutes in mitosis arrested regardless of the number of days/cell cycles in PLK4i. By contrast, the frequency of arrest for daughters of cells that experienced a sub-threshold extension of mitosis (60-90 min) was significantly higher after 2 or 3 compared to 1 day in PLK4i, supporting the conclusion that experiencing sequential sub-threshold prolonged mitoses leads to arrest. (**D**) (*left*) Schematic of p21-mNG intensity measurement in G1, and (right) plot of p21-mNG fluorescence intensity for the indicated conditions. Control is cells treated for 1d with DMSO vehicle. p21-mNG intensity was measured 30 min after mitotic exit. Red dashed line is at the mean value for 2d treatment, when all daughters had robust p21-mNG signal. Error bars are the 95% CI. In the daughters of PLK4i-treated cells experiencing a 60-90 minute sub-threshold mitosis, p21 G1 expression was elevated to a significantly higher degree after successive mitoses (2d) compared to a single mitosis (1d). (**E**) Representative traces of p21-mNG fluorescence intensity versus time after mitotic exit for the indicated conditions. (**F**) (*top*) Outline of experiment: RPE1 FUCCI cells were treated with PLK4i, or DMSO vehicle as a control, for ~3 cell cycles to allow for centrosome depletion and were then filmed for 4 days. (*middle*) Stills from time-lapse sequences. Times listed above panels are in hours after the start of filming. Scale bar is 50 μm. (*bottom*) Graph plotting total cell number versus time. The percentage of cells in G1 phase for the PLK4i-treated culture at the end of filming (96h) is indicated; for the DMSO-treated control, the percentage of cells in G1 phase is shown for the cell density equivalent to that of the PLK4i-treated culture at 96h (indicated by the dashed line). These results indicate that the proliferation arrest observed after treatment with PLK4i is in the G1 phase of the cell cycle.

**Fig. S3.**
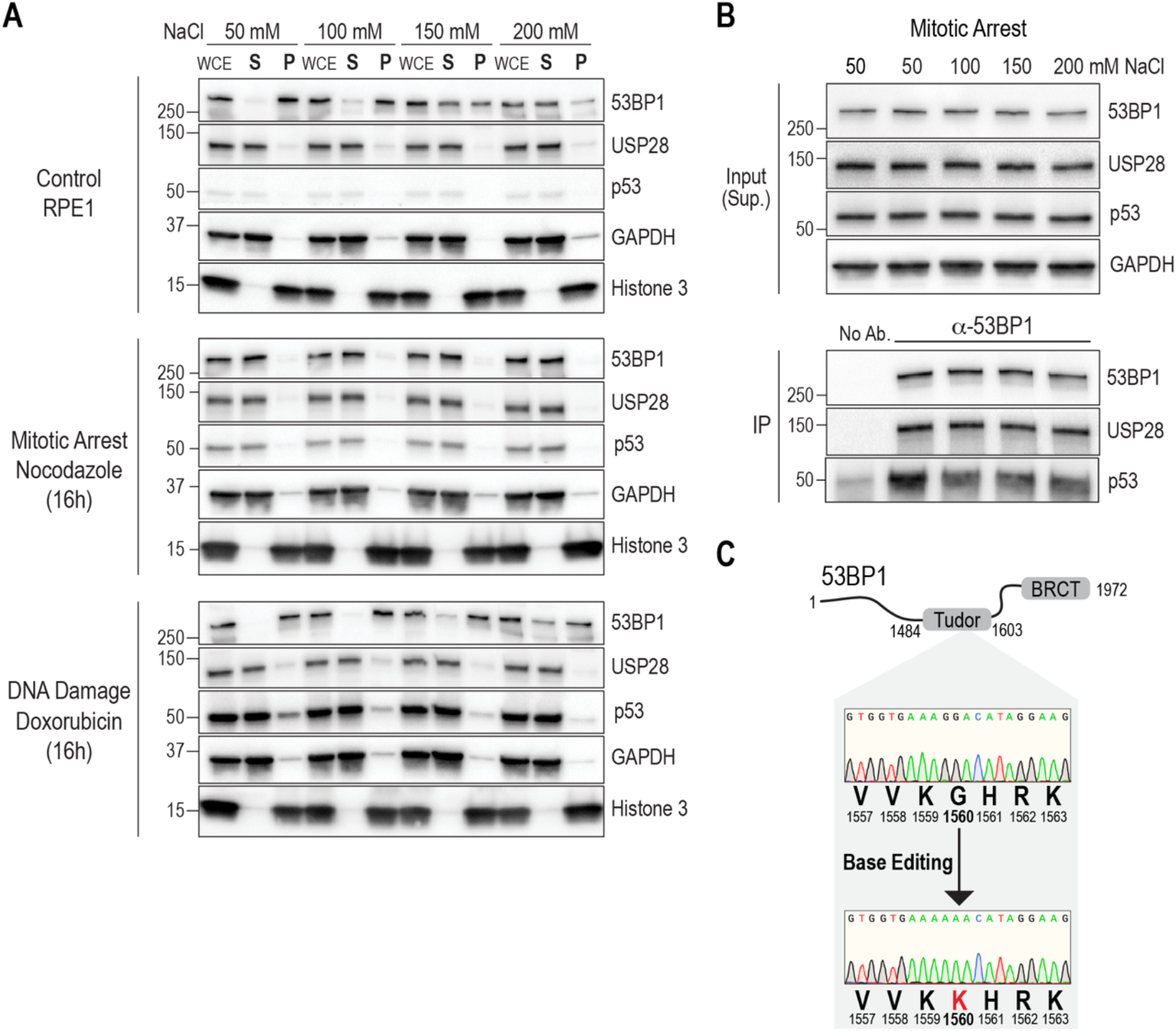
Biochemical analysis of stopwatch components and genotyping of base-edited RPE1 cells. (**A**) Immunoblots of extracts prepared at varying salt concentrations for the indicated treatment conditions. WCE: Whole Cell Extract; S: Supernatant; P: Pellet. GAPDH and histone H3 serve as soluble and chromatin-bound insoluble components, respectively. See Methods for details on the treatment conditions. (**B**) 53BP1 immunoprecipitation from soluble mitotic extracts prepared at different salt concentrations. GAPDH serves as a loading control. (**C**) Sequencing traces of control and base-edited RPE1 cell lines showing mutation of the codon encoding Gly1560 to a codon encoding Lys in both alleles of TP53BP1.

**Fig. S4.**
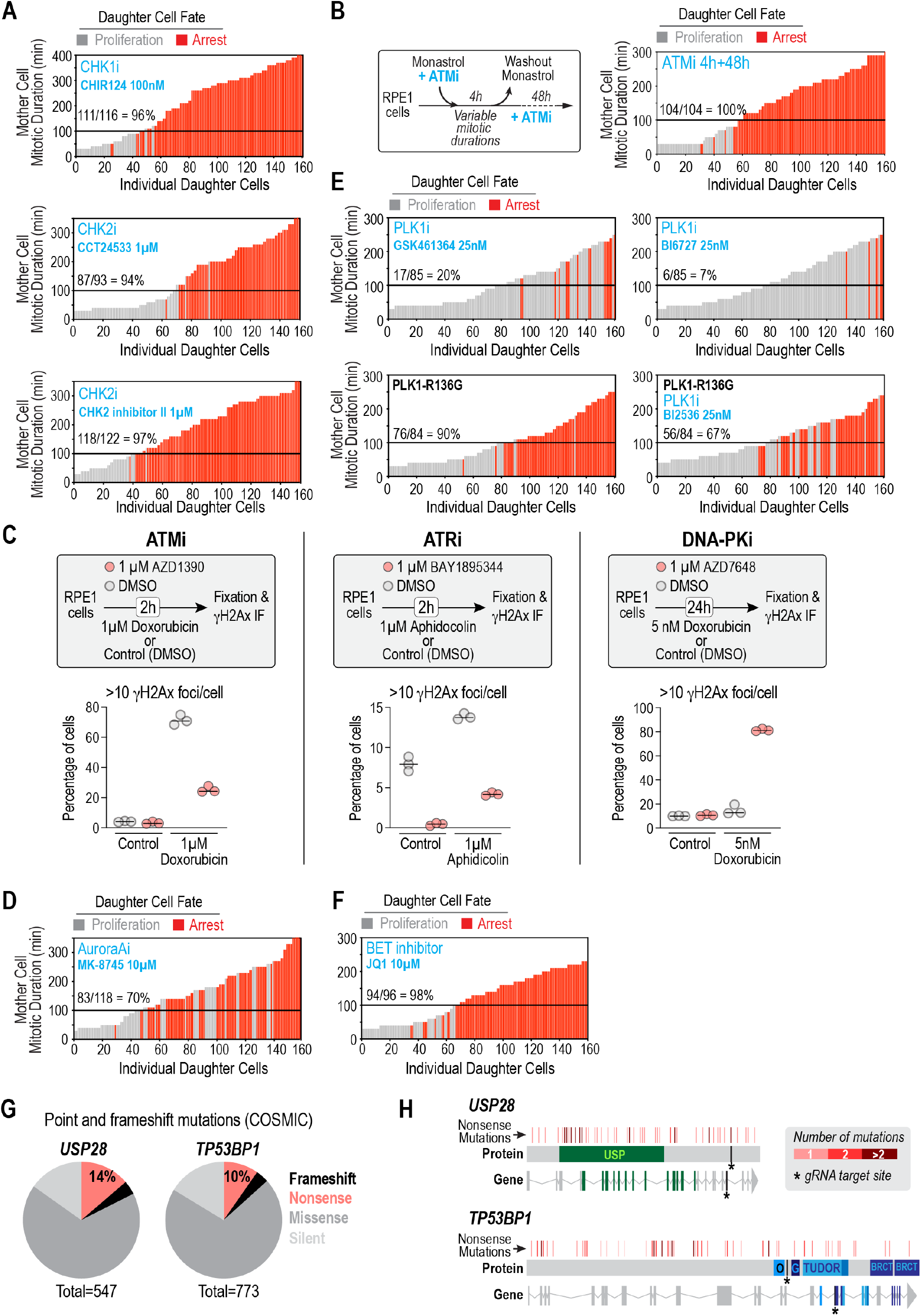
Mitotic stopwatch assays in the presence of inhibitors, assessment of ATM, ATR and DNA-PK inhibitor efficacy, and cancer-associated mutations in *USP28* and *TP53BP1*. (**A**) Mitotic stopwatch assays in RPE1 cells treated with indicated CHK1 and CHK2 inhibitors during the monastrol-induced transient mitotic arrest. CHK1/CHK2 inhibition did not affect stopwatch activity. (**B**) Mitotic stopwatch assay in which ATMi was present prior to and after monastrol washout. Continuous ATM inhibition did not affect stopwatch activity. (**C**) (*top*) Assay schematics and (*bottom*) results of γH2AX immunofluorescence analysis for ATMi, ATRi and DNA-PKi. Three replicates and their mean are plotted; the results indicate that inhibitors were effective in significantly suppressing target activities. (**D**) Mitotic stopwatch assay with the Aurora A-specific inhibitor MK-8475 (52), which led to a mild reduction in stopwatch penetrance. (**E**) (*top*) Mitotic stopwatch assays comparing two different classes of PLK1i; the BI2536 derivative BI6727 (Volasertib; (53)) and GSK461364 which, unlike BI2536 and BI6727, is specific for PLK1 (54). (bottom) comparison of RPE1 cells in which Arg136 of PLK1 was mutated to Gly to induce partial resistance to BI2536. While treatment of control RPE1 cells with 25 nM BI2536 eliminated mitotic stopwatch activity, the same treatment in RPE1-R136G cells was associated with significant stopwatch activity. (**F**) Mitotic stopwatch assay with the BET bromodomain inhibitor JQ1 (55); BI2536 is known to inhibit BET bromodomain proteins with similar potency as JQ1 (56). BET bromodomain inhibition had no effect on stopwatch activity. (**G**) Pie chart showing frequency of frameshift, nonsense, missense, and silent point mutations in the protein-coding regions of USP28 and TP53BP1. These values were extracted from the Catalogue of Somatic Mutations in Cancer (COSMIC, https://cancer.sanger.ac.uk/cosmic). For a gene with no effect on fitness, ~4% of random point mutations are expected to introduce a premature stop codon (57). The significantly higher frequency of nonsense mutations in USP28 and TP53BP1 are consistent with the proposal that they exert a tumor-suppressive function (see also (24)). (**H**) Schematics showing the location of nonsense mutations observed in tumors. Vertical lines are drawn for every 10 amino acids of the protein. Lines are white and invisible unless nonsense mutations were detected, in which case the color of the line indicates the number of detected nonsense mutations. Nonsense mutations are enriched at the beginning of the ubiquitin-specific peptidase (USP) domain of USP28. The asterisks mark sites targeted for introduction of frameshift mutations in non-transformed RPE1 cells.

**Fig. S5.**
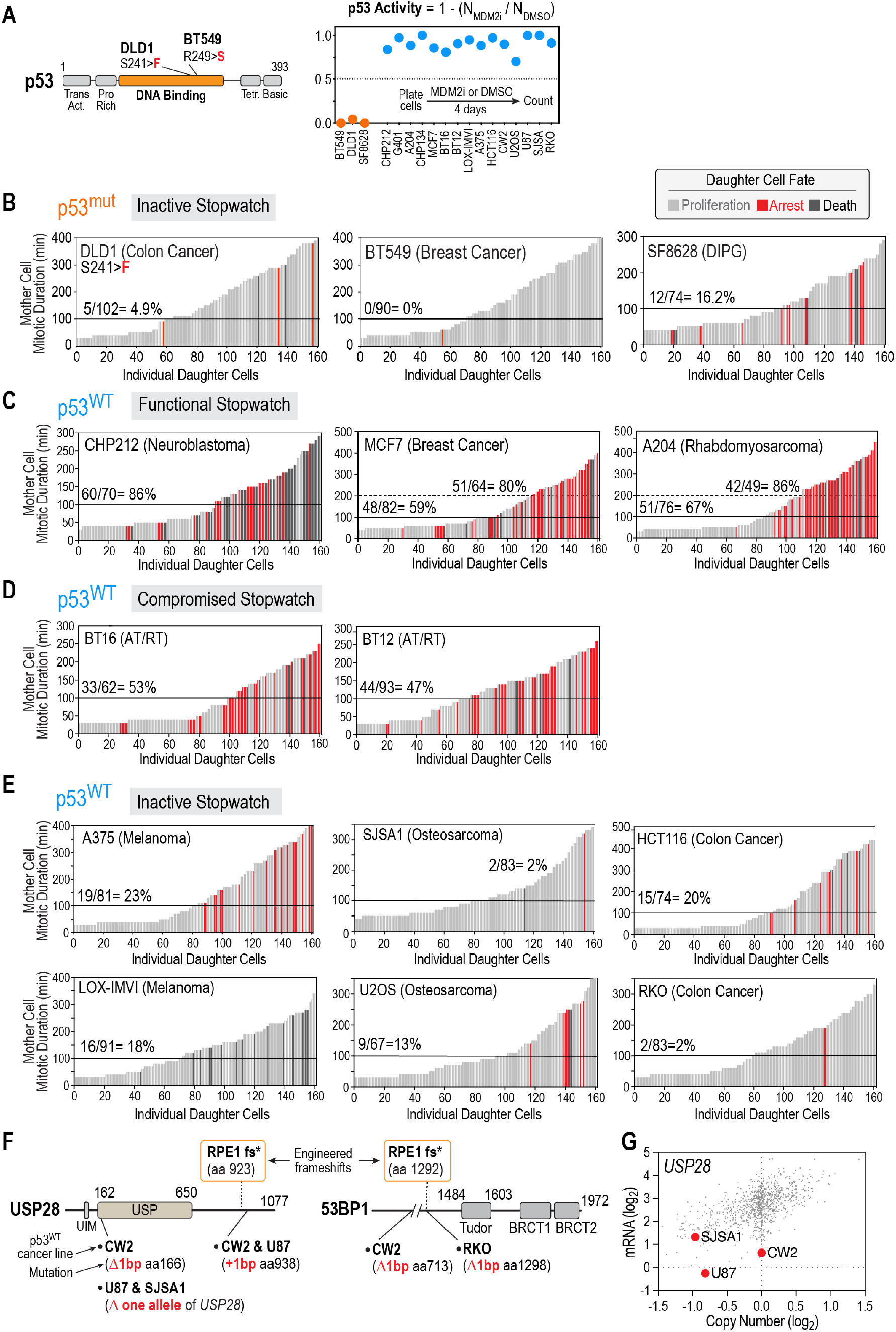
Mitotic stopwatch in p53-wildtype and p53-mutant cancer-derived lines. (**A**) (*left*)Schematic illustrating the location of the cancer-associated p53 mutations in the two indicated cell lines. The precise p53 mutation in SF8628 cells, derived from a pediatric cancer, is not known. (*right*) Experimental confirmation of p53 status by monitoring response to Mdm2i treatment in the cell lines described in *Fig. 4A*. (**B**-**E**) Functional analysis of the mitotic stopwatch in p53-mutant (B) and p53-wildtype (C-E) cancer-derived lines. p53-wildtype cell lines are group based on whether they had a functional (C), compromised (D), or inactive (E) stopwatch. (**F**) Schematics of USP28 and 53BP1 depicting the location of mutations present in the indicated cancer cell lines. The locations targeted to model frameshift mutations in one or both alleles in non-transformed RPE1 cells are also indicated on top. In 4 of the 8 cell lines lacking stopwatch function, mutations were present in USP28 and/or TP53BP1. Immunoblotting revealed bi-allelic mutations and complete loss of USP28 protein in the U87 glioblastoma and CW2 colorectal cancer cell lines (*Fig. 4C*); CW2 also harbored a mutation in one allele of TP53BP1. (**G**) USP28 mRNA level and copy number in the DepMap portal (https://depmap.org/portal/; (58)). Three p53-wildtype cancer lines with an inactive mitotic stopwatch are marked. SJSA1 and U87 both have reduced USP28 copy number, consistent with loss of one allele. CW2 has low mRNA expression and is annotated as having point mutations in both alleles of USP28. Low USP28 mRNA in U87 and CW2 is consistent with loss of USP28 protein in these cell lines (*Fig. 4C*).

**Fig. S6.**
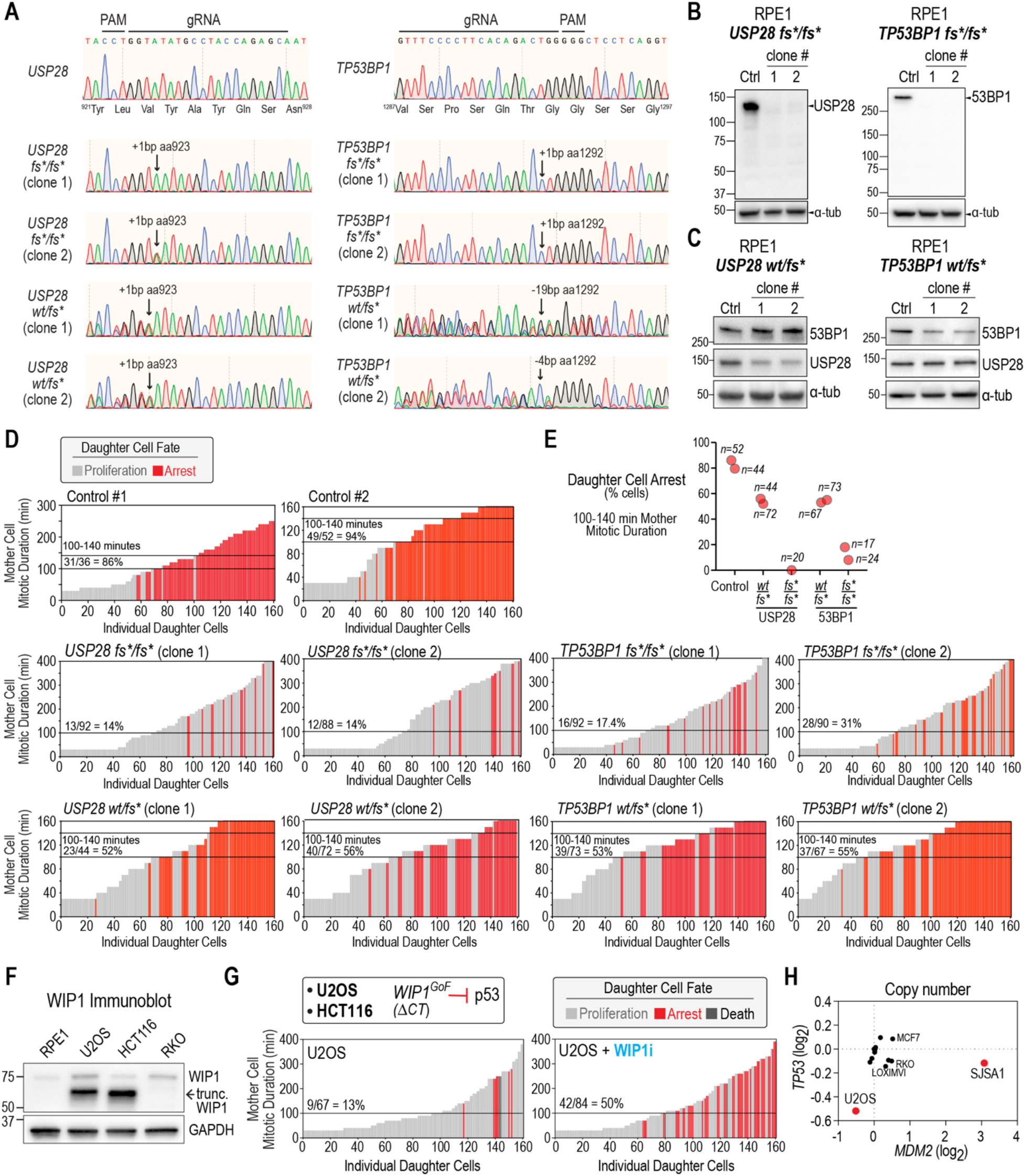
Analysis of mechanisms inactivating stopwatch function in p53-wildtype cancers. (**A**) Genotyping data for frameshift mutations engineered in USP28 and TP53BP1 in RPE1 cells, which were modeled on mutations found in cancers. The PAM and gRNA sequences are indicated on top; sequencing direction was from right to left. (**B, C**) Immunoblot analysis of two independent RPE1 clones with frameshift mutations in either both alleles (B) or only one allele (C) of USP28 or TP53BP1. a-tubulin serves as a loading control. The USP28 antibody was raised against aa 120-160 and the 53BP1 antibody against aa 350-400; both regions are N-terminal to the introduced frameshifts, depicted in *Fig. S5F*. Thus, the absence of a lower molecular weight band is not due to loss of epitopes detected by the antibodies. Introduction of the frameshift in only one allele of either USP28 or TP53BP1 resulted in ~50% protein expression. (**D**) Mitotic stopwatch assays of the homozygous (fs*/fs*) and heterozygous (wt/fs*) frameshift mutant RPE1 cell lines. (**E**) Frequency of arrest of daughter cells produced by mothers with 100-140 min mitosis. Data from two independent clones is shown, except for USP28 fs*/fs*, where only one clone is shown. The reduction in daughter cell arrest frequency for mothers experiencing mitotic durations just above the threshold in the heterozygous clones indicates reduced functionality of the mitotic stopwatch, presumably due to the reduced levels of stopwatch complex components (*Fig. S6C*). (**F**) Immunoblot of the p53-antagonizing phosphatase WIP1 in the indicated cell lines. A highly expressed, C-terminally truncated form of WIP1 is present in both U2OS and HCT116 cells (29). (**G**) (*top*) Schematic depiction of p53 activity suppression by hyperactive, C-terminally-truncated WIP1 phosphatase (29). (*bottom*) Mitotic stopwatch assays in U2OS cells in the absence and presence of a WIP1 inhibitor (10 μM GSK2830371). The WIP1 inhibitor was present during the monastrol treatment used to variably extend mother cell mitosis, and after monastrol washout. The U2OS data in the absence of inhibitor is reproduced from Fig. S5E for comparison. WIP1 inhibition restored significant stopwatch function in U2OS cells. (**H**) Copy number plot for TP53 and MDM2 in a subset of cancer cell lines. SJSA1 has significant amplification of the p53-degrading ubiquitin ligase MDM2 whereas U2OS has reduced copy number of TP53; SJSA1 has also lost one allele of USP28 (*Fig. S5G*). Collectively, the results shown in this figure indicate that direct mutation of core mitotic stopwatch components USP28 and 53BP1 as well as cancer-associated genetic changes that dampen p53 signaling both contribute to compromising mitotic stopwatch function in p53-wildtype cancers.

**Fig. S7.**
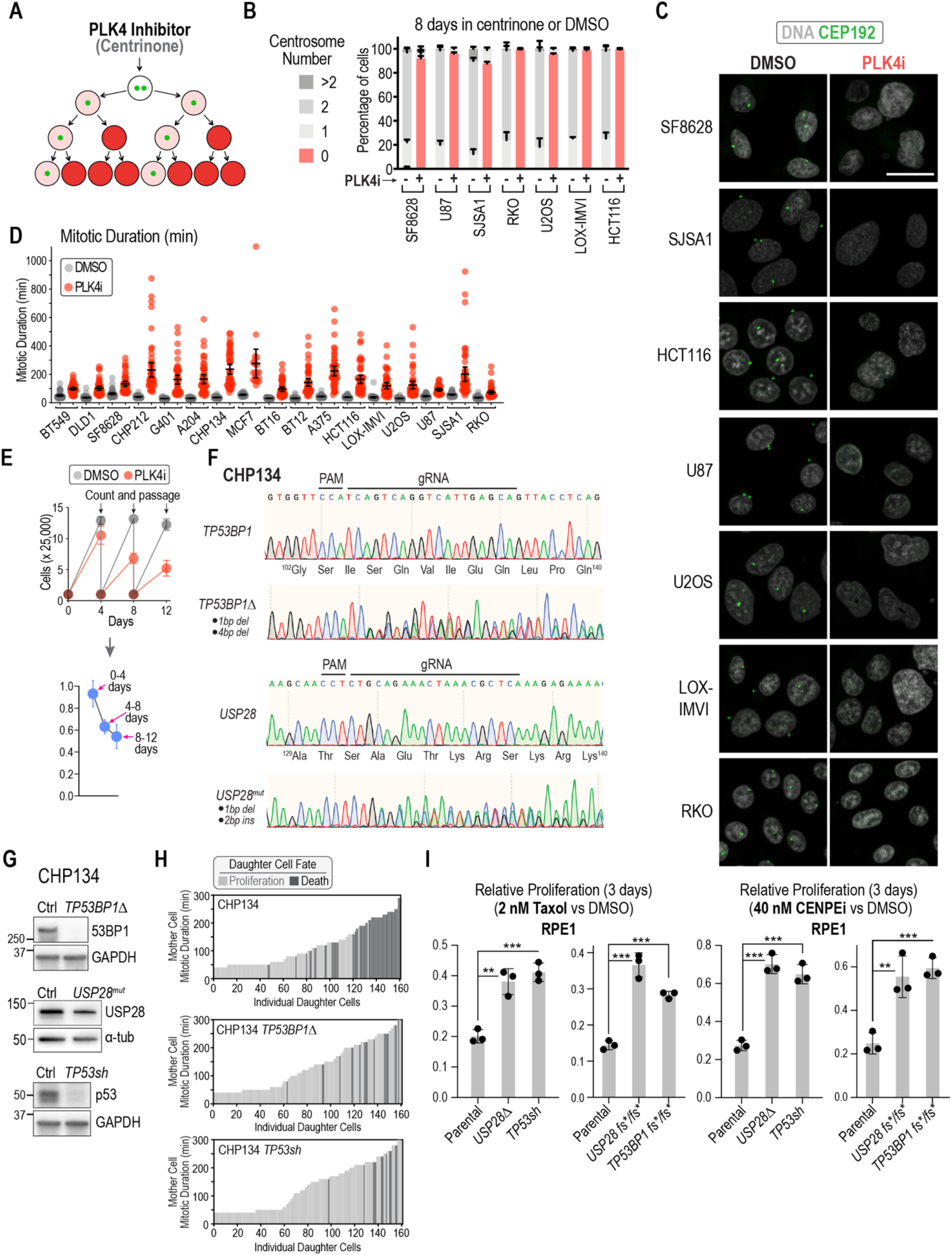
Analysis of PLK4i efficacy in cancer cell lines, characterization of *TP53BP1* and *USP28* mutants generated in the CHP134 neuroblastoma cell line, and sensitivity of RPE1 frameshift mutants to Taxol and CENPEi. (**A**) Treatment with the PLK4i centrinone prevents assembly of new centrosomes, leading to centrosome depletion and the generation of acentrosomal cells as shown in the schematic. (**B**) Graph of centrosome number, measured after 8-day PLK4i or control (DMSO) treatment. (**C**) Images showing centrosome loss, visualized by labeling of the centrosomal component CEP192, in the indicated cancer cell lines. Scale bar, 25 μm. (**D**) Graph plotting mitotic duration measured by live imaging of p53-mutant and p53-wildtype cancer cell lines expressing H2B-RFP. Cells were first treated for 3 cell cycles with PLK4i to induce centrosome loss, or DMSO as a control, prior to live imaging. n=50 for all conditions, except U87 (DMSO, n=42; PLK4i, n=36) and MCF7 (DMSO, n=29; PLK4i, n=19). Error bars are the 95% confidence interval. In all analyzed cell lines, PLK4i treatment resulted in a highly significant increase in mitotic duration (p<0.0001 from a t-test). (**E**) Schematic showing how passaging assay data for DMSO and PLK4i treated cells was converted to the relative proliferation data shown in *Fig. 4D,E*. (**F**) Sequence traces indicating genotypes of mutants generated in the CHP134 neuroblastoma cell line. (**G**) Immunoblots of engineered CHP134 lines. a-tubulin serves as a loading control. As a band of the same molecular weight as USP28 was observed in multiple mutant clones, albeit at reduced levels, we refer to the analyzed cell line as USP28mut. (**H**) Mitotic stopwatch assays of TP53BP1Δ and TP53sh cell lines generated in the CHP134 background. The parental CHP134 line plot from *Fig. 4A* is shown for comparison. (**I**) Mean relative proliferation of parental RPE1 cells and derived USP28Δ, TP53sh, USP28 fs*/fs* and TP53BP1 fs*/fs* clonal lines in 2 nM Taxol (left) or 40 nM CENPEi (right). Error bars are the SD. p-values are from t-tests (**:p<0.01; ***: p<0.001).

**Table S1.** Human cell lines used in this study.

| **Parental cell lines** |  |  |  |  |
| --- | --- | --- | --- | --- |
| **OD cell line code** | **Name** | **Source** | **Clonal or Polyclonal** | **Catalog number** |
| RCL001 | hTERT RPE-1 | ATCC | n/a | CRL-4000 |
| RCL004 | U2OS | ATCC | n/a | HTB-96 |
| RCL006 | HCT116 | ATCC | n/a | CCL-247 |
| RCL008 | CHP212 | ATCC | n/a | CRL-2273 |
| RCL011 | G401 | ATCC | n/a | CRL-1441 |
| RCL012 | SJSA1 | ATCC | n/a | CRL-2098 |
| RCL013 | RKO | ATCC | n/a | CRL-2577 |
| RCL014 | BT12 | Gupta Lab (UCSF) | n/a |  |
| RCL015 | BT16 | Gupta Lab (UCSF) | n/a |  |
| RCL016 | SF8628 | Gupta Lab (UCSF) | n/a |  |
| RCL017 | LOX-IMVI | NCI-60 | n/a |  |
| RCL018 | U87 | ATCC | n/a | HTB-14 |
| RCL019 | CHP134 | Sigma (ECACC) | n/a | 06122002 |
| RCL022 | MCF7 | ATCC | n/a | HTB-22 |
| RCL024 | BT549 | ATCC | n/a | HTB-122 |
| RCL043 | DLD1 | Sigma (ECACC) | n/a | 90102540 |
| RCL045 | A204 | ATCC | n/a | HTB-82 |
| RCL060 | A375 | ATCC | n/a | CRL-1619 |
| RCL061 | CW2 | Cell Bank Riken | n/a | RCB0778 |
| **Engineered cell lines** |  |  |  |  |
| **OD cell line code** | **Parental line** | **Modification(s)** | **Clonal or Polyclonal** | **Reference** |
| ODCL0002 | hTERT RPE-1 | *USP28Δ* | Clonal | Meitinger et al. 2016 |
| ODCL0035 | hTERT RPE-1 | *EF-1 $\alpha^{pro}$-H2B-mRFP* | Polyclonal | Meitinger et al. 2016 |
| ODCL0036 | CHP212 | *EF-1 $\alpha^{pro}$-H2B-mRFP* | Polyclonal | Meitinger et al. 2020 |
| ODCL0039 | G401 | *EF-1 $\alpha^{pro}$-H2B-mRFP* | Polyclonal | This study |
| ODCL0040 | U2OS | *EF-1 $\alpha^{pro}$-H2B-mRFP* | Polyclonal | This study |
| ODCL0041 | SJSA1 | *EF-1 $\alpha^{pro}$-H2B-mRFP* | Polyclonal | This study |
| ODCL0042 | RKO | *EF-1 $\alpha^{pro}$-H2B-mRFP* | Polyclonal | This study |
| ODCL0043 | HCT116 | *EF-1 $\alpha^{pro}$-H2B-mRFP* | Polyclonal | This study |
| ODCL0044 | BT12 | *EF-1 $\alpha^{pro}$-H2B-mRFP* | Polyclonal | This study |
| ODCL0045 | BT16 | *EF-1 $\alpha^{pro}$-H2B-mRFP* | Polyclonal | This study |
| ODCL0046 | SF8628 | *EF-1 $\alpha^{pro}$-H2B-mRFP* | Polyclonal | This study |
| ODCL0047 | LOX-IMVI | *EF-1 $\alpha^{pro}$-H2B-mRFP* | Polyclonal | This study |
| ODCL0048 | U87 | *EF-1 $\alpha^{pro}$-H2B-mRFP* | Polyclonal | This study |
| ODCL0049 | CHP134 | *EF-1 $\alpha^{pro}$-H2B-mRFP* | Polyclonal | Meitinger et al. 2020 |
| ODCL0050 | CHP134 | *TP53BP1Δ* | Clonal | This study |
| ODCL0051 | CHP134 | *TP53BP1Δ, EF-1$\alpha^{pro}$-H2B-mRFP* | Polyclonal | This study |
| ODCL0052 | CHP134 | *TP53-sh* | Polyclonal | This study |
| ODCL0053 | CHP134 | *TP53-sh, EF-1$\alpha^{pro}$-H2B-mRFP* | Polyclonal | This study |
| ODCL0054 | hTERT RPE-1 | *TP53-sh* | Polyclonal | Meitinger et al. 2016 |
| ODCL0059 | hTERT RPE-1 | *CDKN1A(p21)Δ* | Clonal | This study |
| ODCL0077 | hTERT RPE-1 | *CEP192-mNeonGreen USP28Δ TRE3G$^{pro}$-Cas9* | Clonal | Meitinger et al. 2020 |
| ODCL0130 | MCF7 | *EF-1$\alpha^{pro}$-H2B-mRFP* | Polyclonal | This study |
| ODCL0131 | A204 | *EF-1$\alpha^{pro}$-H2B-mRFP* | Polyclonal | This study |
| ODCL0132 | CW2 | *EF-1$\alpha^{pro}$-H2B-mRFP* | Polyclonal | This study |
| ODCL0133 | hTERT RPE-1 | *TP53-sh EF-1$\alpha^{pro}$-H2B-mRFP* | Polyclonal | Meitinger et al. 2016 |
| ODCL0191 | hTERT RPE-1 | *CEP192-mNeonGreen TRE3G$^{pro}$-Cas9* | Clonal | This study |
| ODCL0192 | hTERT RPE-1 | *CEP192-mNeonGreen TRE3G$^{pro}$-Cas9 TP53-sh* | Polyclonal | This study |
| ODCL0247 | hTERT RPE-1 | *TP53BP1Δ* | Clonal | Meitinger et al. 2016 |
| ODCL0248 | hTERT RPE-1 | *TP53BP1Δ H2B-RFP* | Polyclonal | Meitinger et al. 2016 |
| ODCL0249 | hTERT RPE-1 | *USP28Δ H2B-RFP* | Polyclonal | Meitinger et al. 2016 |
| ODCL0250 | hTERT RPE-1 | *mVenus-hGeminin1-110 EF-1$\alpha^{pro}$-H2B-mRFP* | Polyclonal | This study |
| ODCL0251 | hTERT RPE-1 | *CDKN1A(p21)Δ EF-1$\alpha^{pro}$-H2B-mRFP* | Polyclonal | This study |
| ODCL0254 | hTERT RPE-1 | *EF-1$\alpha^{pro}$-APOBEC-1-Cas9(D10A)-UGI; U6$^{pro}$-gRNA-TP53BP1(G1560K)* | Clonal | This study |
| ODCL0258 | hTERT RPE-1 | *TP53BP1 wt/fs\* (19bp del)* | Clone 1 | This study |
| ODCL0259 | hTERT RPE-1 | *TP53BP1 wt/fs\* (4bp del)* | Clone 2 | This study |
| ODCL0261 | hTERT RPE-1 | *EF-1$\alpha^{pro}$-APOBEC-1-Cas9(D10A)-UGI; U6$^{pro}$-gRNA-TP53BP1(G1560K); EF-1$\alpha^{pro}$-H2B-mRFP* | Polyclonal | This study |
| ODCL0262 | hTERT RPE-1 | *EF-1$\alpha^{pro}$-APOBEC-1-Cas9(D10A)-UGI; U6$^{pro}$-gRNA-TP53BP1(G1560K); EF-1$\alpha^{pro}$-H2B-mRFP* | Polyclonal | This study |
| ODCL0263 | hTERT RPE-1 | *TP53BP1 wt/fs\* (19bp del) EF-1$\alpha^{pro}$-H2B-mRFP* | Polyclonal | This study |
| ODCL0264 | hTERT RPE-1 | *TP53BP1 wt/fs\* (4bp del) EF-1 $\alpha^{pro}$-H2B-mRFP* | Polyclonal | This study |
| ODCL0330 | CHP134 | *USP28Δ* | Clonal | This study |
| ODCL0340 | CHP134 | *USP28Δ, EF-1 $\alpha^{pro}$-H2B-mRFP* | Polyclonal | This study |
| ODCL0358 | DLD1 | *EF-1 $\alpha^{pro}$-H2B-mRFP* | Polyclonal | This study |
| ODCL0359 | BT549 | *EF-1 $\alpha^{pro}$-H2B-mRFP* | Polyclonal | This study |
| ODCL0365 | hTERT RPE-1 | *USP28 wt/fs\* (1bp ins)* | Clone 1 | This study |
| ODCL0368 | hTERT RPE-1 | *USP28 wt/fs\* (1bp ins)* | Clone 2 | This study |
| ODCL0369 | hTERT RPE-1 | *USP28 wt/fs\* (1bp ins) EF-1 $\alpha^{pro}$-H2B-mRFP* | Polyclonal | This study |
| ODCL0371 | hTERT RPE-1 | *USP28 wt/fs\* (1bp ins) EF-1 $\alpha^{pro}$-H2B-mRFP* | Polyclonal | This study |
| ODCL0374 | hTERT RPE-1 | *CEP192-mNeonGreen TRE3G<sup>pro</sup>-Cas9 TP53-sh U6<sup>pro</sup>-gRNA-Comet EF-1 $\alpha^{pro}$-H2B-mRFP* | Polyclonal | This study |
| ODCL0375 | hTERT RPE-1 | *CEP192-mNeonGreen USP28Δ TRE3G<sup>pro</sup>-Cas9 U6<sup>pro</sup>-gRNA-Comet EF-1 $\alpha^{pro}$-H2B-mRFP* | Polyclonal | This study |
| ODCL0381 | hTERT RPE-1 | *USP28 fs\*/fs\* (1bp ins)* | Clone 1 | This study |
| ODCL0382 | hTERT RPE-1 | *USP28 fs\*/fs\* (1bp ins)* | Clone 2 | This study |
| ODCL0384 | hTERT RPE-1 | *USP28 fs\*/fs\* (1bp ins) EF-1 $\alpha^{pro}$-H2B-mRFP* | Polyclonal | This study |
| ODCL0385 | hTERT RPE-1 | *USP28 fs\*/fs\* (1bp ins) EF-1 $\alpha^{pro}$-H2B-mRFP* | Polyclonal | This study |
| ODCL0393 | A375 | *EF-1 $\alpha^{pro}$-H2B-mRFP* | Polyclonal | This study |
| ODCL0404 | hTERT RPE-1 | *P21-NeonGreen* | Clonal | This study |
| ODCL0405 | hTERT RPE-1 | *P21-NeonGreen EF-1 $\alpha^{pro}$-H2B-mRFP* | Polyclonal | This study |
| ODCL0406 | hTERT RPE-1 | *TP53BP1 fs\*/fs\* (1bp ins) EF-1 $\alpha^{pro}$-H2B-mRFP* | Polyclonal | This study |
| ODCL0407 | hTERT RPE-1 | *TP53BP1 fs\*/fs\* (1bp ins) EF-1 $\alpha^{pro}$-H2B-mRFP* | Polyclonal | This study |
| ODCL0408 | hTERT RPE-1 | *CEP192-mNeonGreen USP28Δ TRE3G<sup>pro</sup>-Cas9 U6<sup>pro</sup>-gRNA-Comet* | Polyclonal | This study |
| ODCL0409 | hTERT RPE-1 | *CEP192-mNeonGreen TRE3G<sup>pro</sup>-Cas9 U6<sup>pro</sup>-gRNA-Comet* | Polyclonal | This study |
| ODCL0410 | hTERT RPE-1 | *CEP192-mNeonGreen TRE3G<sup>pro</sup>-Cas9 TP53-sh U6<sup>pro</sup>-gRNA-Comet* | Polyclonal | This study |
| ODCL0411 | hTERT RPE-1 | *CEP192-mNeonGreen TRE3G<sup>pro</sup>-Cas9 U6<sup>pro</sup>-* *gRNA-Comet EF-1 $\alpha^{pro}$-H2B-mRFP* | Polyclonal | This study |
| ODCL0412 | hTERT RPE-1 | *TP53BP1 fs\*/fs\* (1bp ins)* | Clone 1 | This study |
| ODCL0413 | hTERT RPE-1 | *TP53BP1 fs\*/fs\* (1bp ins)* | Clone 2 | This study |
| ODCL0416 | hTERT RPE-1 | *CEP192-mNeonGreen*<br>*TRE3G<sup>pro</sup>-Cas9 U6<sup>pro</sup>-*<br>*gRNA-AAVS1* | Polyclonal | This study |
| ODCL0418 | hTERT RPE-1 | *CEP192-mNeonGreen*<br>*TRE3G<sup>pro</sup>-Cas9 U6<sup>pro</sup>-*<br>*gRNA-AAVS1 EF-1 $\alpha^{pro}$-*<br>*H2B-mRFP* | Polyclonal | This study |
| ODCL432 | hTERT RPE-1 | *PLK1-R136G* | Clonal | This study |

(pro, Promoter; del, deletion; ins, insertion)

**Table S2.**
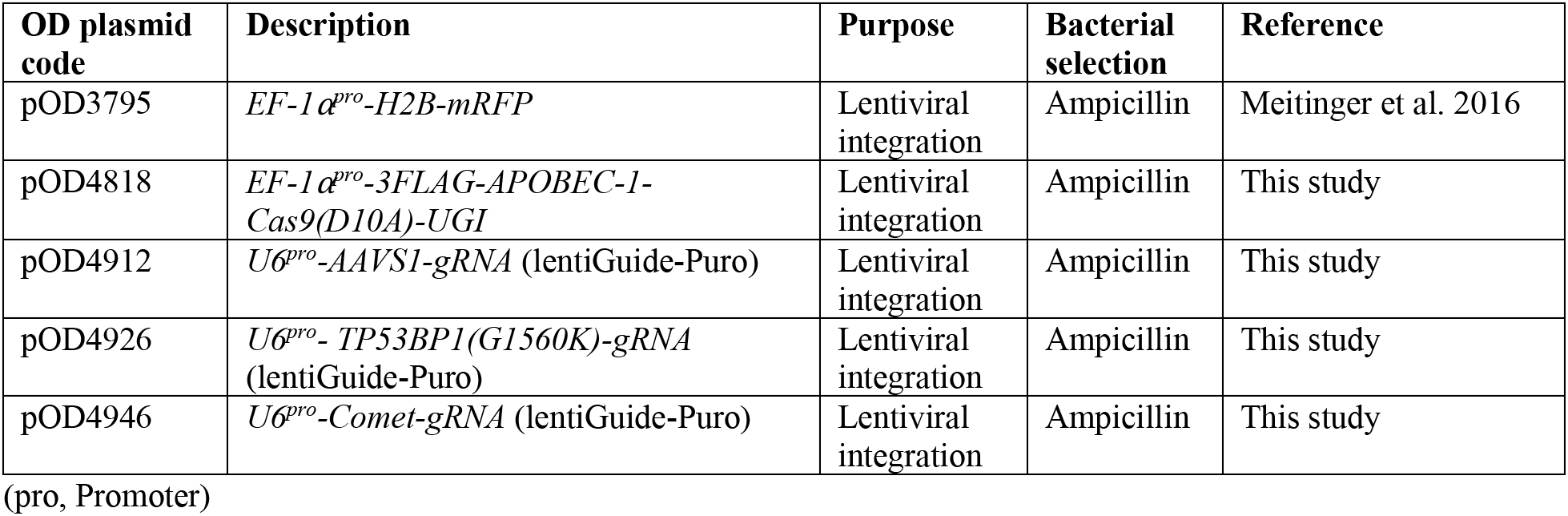
Plasmids used in this study.

| **OD plasmid code** | **Description** | **Purpose** | **Bacterial selection** | **Reference** |
| --- | --- | --- | --- | --- |
| pOD3795 | *EF-1$\alpha^{pro}$-H2B-mRFP* | Lentiviral integration | Ampicillin | Meitinger et al. 2016 |
| pOD4818 | *EF-1$\alpha^{pro}$-3FLAG-APOBEC-1-Cas9(D10A)-UGI* | Lentiviral integration | Ampicillin | This study |
| pOD4912 | *U6$^{pro}$-AAVS1-gRNA* (lentiGuide-Puro) | Lentiviral integration | Ampicillin | This study |
| pOD4926 | *U6$^{pro}$-TP53BP1(G1560K)-gRNA* (lentiGuide-Puro) | Lentiviral integration | Ampicillin | This study |
| pOD4946 | *U6$^{pro}$-Comet-gRNA* (lentiGuide-Puro) | Lentiviral integration | Ampicillin | This study |

(pro, Promoter)

## References

Araujo, A.R., L. Gelens, R.S. Sheriff, and S.D. Santos. 2016. Positive Feedback Keeps Duration of Mitosis Temporally Insulated from Upstream Cell-Cycle Events. Mol Cell. 64:362–375.

Barr, A.R., S. Cooper, F.S. Heldt, F. Butera, H. Stoy, J. Mansfeld, B. Novak, and C. Bakal. 2017. DNA damage during S-phase mediates the proliferation-quiescence decision in the subsequent G1 via p21 expression. Nat Commun. 8:14728.

Bazzi, H., and K.V. Anderson. 2014. Acentriolar mitosis activates a p53-dependent apoptosis pathway in the mouse embryo. Proc Natl Acad Sci U S A. 111:E1491–1500.

Bernabeu, E., M. Cagel, E. Lagomarsino, M. Moretton, and D.A. Chiappetta. 2017. Paclitaxel: What has been done and the challenges remain ahead. Int J Pharm. 526:474–495.

Brinkman, E.K., T. Chen, M. Amendola, and B. van Steensel. 2014. Easy quantitative assessment of genome editing by sequence trace decomposition. Nucleic Acids Res. 42:e168.

Cimini, D., B. Howell, P. Maddox, A. Khodjakov, F. Degrassi, and E.D. Salmon. 2001. Merotelic kinetochore orientation is a major mechanism of aneuploidy in mitotic mammalian tissue cells. J Cell Biol. 153:517–527.

Corbett, K.D. 2017. Molecular Mechanisms of Spindle Assembly Checkpoint Activation and Silencing. Prog Mol Subcell Biol. 56:429–455.

Cuella-Martin, R., S.B. Hayward, X. Fan, X. Chen, J.W. Huang, A. Taglialatela, G. Leuzzi, J. Zhao, R. Rabadan, C. Lu, Y. Shen, and A. Ciccia. 2021. Functional interrogation of DNA damage response variants with base editing screens. Cell. 184:1081–1097 e1019.

Davoli, T., A.W. Xu, K.E. Mengwasser, L.M. Sack, J.C. Yoon, P.J. Park, and S.J. Elledge. 2013. Cumulative haploinsufficiency and triplosensitivity drive aneuploidy patterns and shape the cancer genome. Cell. 155:948–962.

Derbyshire, D.J., B.P. Basu, L.C. Serpell, W.S. Joo, T. Date, K. Iwabuchi, and A.J. Doherty. 2002. Crystal structure of human 53BP1 BRCT domains bound to p53 tumour suppressor. EMBO J. 21:3863–3872.

Dumontet, C., and M.A. Jordan. 2010. Microtubule-binding agents: a dynamic field of cancer therapeutics. Nat Rev Drug Discov. 9:790–803.

Durant, S.T., L. Zheng, Y. Wang, K. Chen, L. Zhang, T. Zhang, Z. Yang, L. Riches, A.G. Trinidad, J.H.L. Fok, T. Hunt, K.G. Pike, J. Wilson, A. Smith, N. Colclough, V.P. Reddy, A. Sykes, A. Janefeldt, P. Johnstrom, K. Varnas, A. Takano, S. Ling, J. Orme, J. Stott, C. Roberts, I. Barrett, G. Jones, M. Roudier, A. Pierce, J. Allen, J. Kahn, A. Sule, J. Karlin, A. Cronin, M. Chapman, K. Valerie, R. Illingworth, and M. Pass. 2018. The brain-penetrant clinical ATM inhibitor AZD1390 radiosensitizes and improves survival of preclinical brain tumor models. Sci Adv. 4:eaat1719.

Fernandez-Vidal, A., J. Vignard, and G. Mirey. 2017. Around and beyond 53BP1 Nuclear Bodies. Int J Mol Sci. 18.

Ferrell, J.E., Jr., and S.H. Ha. 2014. Ultrasensitivity part II: multisite phosphorylation, stoichiometric inhibitors, and positive feedback. Trends Biochem Sci. 39:556–569.

Fok, J.H.L., A. Ramos-Montoya, M. Vazquez-Chantada, P.W.G. Wijnhoven, V. Follia, N. James, P.M. Farrington, A. Karmokar, S.E. Willis, J. Cairns, J. Nikkila, D. Beattie, G.M. Lamont, M.R.V. Finlay, J. Wilson, A. Smith, L.O. O’Connor, S. Ling, S.E. Fawell, M.J. O’Connor, S.J. Hollingsworth, E. Dean, F.W. Goldberg, B.R. Davies, and E.B. Cadogan. 2019. AZD7648 is a potent and selective DNA-PK inhibitor that enhances radiation, chemotherapy and olaparib activity. Nat Commun. 10:5065.

Fong, C.S., G. Mazo, T. Das, J. Goodman, M. Kim, B.P. O’Rourke, D. Izquierdo, and M.F. Tsou. 2016. 53BP1 and USP28 mediate p53-dependent cell cycle arrest in response to centrosome loss and prolonged mitosis. Elife. 5.

Garribba, L., and S. Santaguida. 2022. The Dynamic Instability of the Aneuploid Genome. Front Cell Dev Biol. 10:838928.

Gilmartin, A.G., T.H. Faitg, M. Richter, A. Groy, M.A. Seefeld, M.G. Darcy, X. Peng, K. Federowicz, J. Yang, S.Y. Zhang, E. Minthorn, J.P. Jaworski, M. Schaber, S. Martens, D.E. McNulty, R.H. Sinnamon, H. Zhang, R.B. Kirkpatrick, N. Nevins, G. Cui, B. Pietrak, E. Diaz, A. Jones, M. Brandt, B. Schwartz, D.A. Heerding, and R. Kumar. 2014. Allosteric Wip1 phosphatase inhibition through flap-subdomain interaction. Nat Chem Biol. 10:181–187.

Insolera, R., H. Bazzi, W. Shao, K.V. Anderson, and S.H. Shi. 2014. Cortical neurogenesis in the absence of centrioles. Nat Neurosci. 17:1528–1535.

Joo, W.S., P.D. Jeffrey, S.B. Cantor, M.S. Finnin, D.M. Livingston, and N.P. Pavletich. 2002. Structure of the 53BP1 BRCT region bound to p53 and its comparison to the Brca1 BRCT structure. Genes Dev. 16:583–593.

Kaulich, M., and S.F. Dowdy. 2015. Combining CRISPR/Cas9 and rAAV Templates for Efficient Gene Editing. Nucleic Acid Ther. 25:287–296.

Khoo, K.H., C.S. Verma, and D.P. Lane. 2014. Drugging the p53 pathway: understanding the route to clinical efficacy. Nat Rev Drug Discov. 13:217–236.

Kim, D.H., J.S. Han, P. Ly, Q. Ye, M.A. McMahon, K. Myung, K.D. Corbett, and D.W. Cleveland. 2018. TRIP13 and APC15 drive mitotic exit by turnover of interphase-and unattached kinetochore-produced MCC. Nat Commun. 9:4354.

Kleiblova, P., I.A. Shaltiel, J. Benada, J. Sevcik, S. Pechackova, P. Pohlreich, E.E. Voest, P. Dundr, J. Bartek, Z. Kleibl, R.H. Medema, and L. Macurek. 2013. Gain-of-function mutations of PPM1D/Wip1 impair the p53-dependent G1 checkpoint. J Cell Biol. 201:511–521.

Knobel, P.A., R. Belotserkovskaya, Y. Galanty, C.K. Schmidt, S.P. Jackson, and T.H. Stracker. 2014. USP28 is recruited to sites of DNA damage by the tandem BRCT domains of 53BP1 but plays a minor role in double-strand break metabolism. Mol Cell Biol. 34:2062–2074.

Kurose, A., T. Tanaka, X. Huang, F. Traganos, W. Dai, and Z. Darzynkiewicz. 2006. Effects of hydroxyurea and aphidicolin on phosphorylation of ataxia telangiectasia mutated on Ser 1981 and histone H2AX on Ser 139 in relation to cell cycle phase and induction of apoptosis. Cytometry A. 69:212–221.

Lambrus, B.G., V. Daggubati, Y. Uetake, P.M. Scott, K.M. Clutario, G. Sluder, and A.J. Holland. 2016. A USP28-53BP1-p53-p21 signaling axis arrests growth after centrosome loss or prolonged mitosis. J Cell Biol. 214:143–153.

Li, R., and J. Zhu. 2022. Effects of aneuploidy on cell behaviour and function. Nat Rev Mol Cell Biol. 23:250–265.

Marjanovic, M., C. Sanchez-Huertas, B. Terre, R. Gomez, J.F. Scheel, S. Pacheco, P.A. Knobel, A. Martinez-Marchal, S. Aivio, L. Palenzuela, U. Wolfrum, P.J. McKinnon, J.A. Suja, I. Roig, V. Costanzo, J. Luders, and T.H. Stracker. 2015. CEP63 deficiency promotes p53-dependent microcephaly and reveals a role for the centrosome in meiotic recombination. Nat Commun. 6:7676.

Meitinger, F., J.V. Anzola, M. Kaulich, A. Richardson, J.D. Stender, C. Benner, C.K. Glass, S.F. Dowdy, A. Desai, A.K. Shiau, and K. Oegema. 2016. 53BP1 and USP28 mediate p53 activation and G1 arrest after centrosome loss or extended mitotic duration. J Cell Biol. 214:155–166.

Meitinger, F., M. Ohta, K.Y. Lee, S. Watanabe, R.L. Davis, J.V. Anzola, R. Kabeche, D.A. Jenkins, A.K. Shiau, A. Desai, and K. Oegema. 2020. TRIM37 controls cancer-specific vulnerability to PLK4 inhibition. Nature. 585:440–446.

Olivier, M., M. Hollstein, and P. Hainaut. 2010. TP53 mutations in human cancers: origins, consequences, and clinical use. Cold Spring Harb Perspect Biol. 2:a001008.

Penna, L.S., J.A.P. Henriques, and D. Bonatto. 2017. Anti-mitotic agents: Are they emerging molecules for cancer treatment? Pharmacol Ther. 173:67–82.

Phan, T.P., and A.J. Holland. 2021. Time is of the essence: the molecular mechanisms of primary microcephaly. Genes Dev. 35:1551–1578.

Phan, T.P., A.L. Maryniak, C.A. Boatwright, J. Lee, A. Atkins, A. Tijhuis, D.C. Spierings, H. Bazzi, F. Foijer, P.W. Jordan, T.H. Stracker, and A.J. Holland. 2021. Centrosome defects cause microcephaly by activating the 53BP1-USP28-TP53 mitotic surveillance pathway. EMBO J. 40:e106118.

Qian, X., A. McDonald, H.J. Zhou, N.D. Adams, C.A. Parrish, K.J. Duffy, D.M. Fitch, R. Tedesco, L.W. Ashcraft, B. Yao, H. Jiang, J.K. Huang, M.V. Marin, C.E. Aroyan, J. Wang, S. Ahmed, J.L. Burgess, A.M. Chaudhari, C.A. Donatelli, M.G. Darcy, L.H. Ridgers, K.A. Newlander, S.J. Schmidt, D. Chai, M. Colon, M.N. Zimmerman, L. Lad, R. Sakowicz, S. Schauer, L. Belmont, R. Baliga, D.W. Pierce, J.T. Finer, Z. Wang, B.P. Morgan, D.J. Morgans, Jr., K.R. Auger, C.M. Sung, J.D. Carson, L. Luo, E.D. Hugger, R.A. Copeland, D. Sutton, J.D. Elliott, J.R. Jackson, K.W. Wood, D. Dhanak, G. Bergnes, and S.D. Knight. 2010. Discovery of the First Potent and Selective Inhibitor of Centromere-Associated Protein E: GSK923295. ACS Med Chem Lett. 1:30–34.

Ran, F.A., P.D. Hsu, J. Wright, V. Agarwala, D.A. Scott, and F. Zhang. 2013. Genome engineering using the CRISPR-Cas9 system. Nat Protoc. 8:2281–2308.

Sakaue-Sawano, A., T. Kobayashi, K. Ohtawa, and A. Miyawaki. 2011. Drug-induced cell cycle modulation leading to cell-cycle arrest, nuclear mis-segregation, or endoreplication. BMC Cell Biol. 12:2.

Sanjana, N.E., O. Shalem, and F. Zhang. 2014. Improved vectors and genome-wide libraries for CRISPR screening. Nat Methods. 11:783–784.

Schindelin, J., I. Arganda-Carreras, E. Frise, V. Kaynig, M. Longair, T. Pietzsch, S. Preibisch, C. Rueden, S. Saalfeld, B. Schmid, J.Y. Tinevez, D.J. White, V. Hartenstein, K. Eliceiri, P. Tomancak, and A. Cardona. 2012. Fiji: an open-source platform for biological-image analysis. Nat Methods. 9:376–682.

Shoshani, O., S.F. Brunner, R. Yaeger, P. Ly, Y. Nechemia-Arbely, D.H. Kim, R. Fang, G.A. Castillon, M. Yu, J.S.Z. Li, Y. Sun, M.H. Ellisman, B. Ren, P.J. Campbell, and D.W. Cleveland. 2021. Chromothripsis drives the evolution of gene amplification in cancer. Nature. 591:137–141.

Tischer, J., and F. Gergely. 2019. Anti-mitotic therapies in cancer. J Cell Biol. 218:10–11.

Uetake, Y., and G. Sluder. 2010. Prolonged prometaphase blocks daughter cell proliferation despite normal completion of mitosis. Curr Biol. 20:1666–1671.

Umbreit, N.T., C.Z. Zhang, L.D. Lynch, L.J. Blaine, A.M. Cheng, R. Tourdot, L. Sun, H.F. Almubarak, K. Judge, T.J. Mitchell, A. Spektor, and D. Pellman. 2020. Mechanisms generating cancer genome complexity from a single cell division error. Science. 368.

Wong, Y.L., J.V. Anzola, R.L. Davis, M. Yoon, A. Motamedi, A. Kroll, C.P. Seo, J.E. Hsia, S.K. Kim, J.W. Mitchell, B.J. Mitchell, A. Desai, T.C. Gahman, A.K. Shiau, and K. Oegema. 2015. Cell biology. Reversible centriole depletion with an inhibitor of Polo-like kinase 4. Science. 348:1155–1160.

Xiao, C., M. Grzonka, C. Meyer-Gerards, M. Mack, R. Figge, and H. Bazzi. 2021. Gradual centriole maturation associates with the mitotic surveillance pathway in mouse development. EMBO Rep. 22:e51127.

Yang, H.W., M. Chung, T. Kudo, and T. Meyer. 2017. Competing memories of mitogen and p53 signalling control cell-cycle entry. Nature. 549:404–408.

Yeow, Z.Y., B.G. Lambrus, R. Marlow, K.H. Zhan, M.A. Durin, L.T. Evans, P.M. Scott, T. Phan, E. Park, L.A. Ruiz, D. Moralli, E.G. Knight, L.M. Badder, D. Novo, S. Haider, C.M. Green, A.N.J. Tutt, C.J. Lord, J.R. Chapman, and A.J. Holland. 2020. Targeting TRIM37-driven centrosome dysfunction in 17q23-amplified breast cancer. Nature. 585:447–452.

Zafra, M.P., E.M. Schatoff, A. Katti, M. Foronda, M. Breinig, A.Y. Schweitzer, A. Simon, T. Han, S. Goswami, E. Montgomery, J. Thibado, E.R. Kastenhuber, F.J. Sanchez-Rivera, J. Shi, C.R. Vakoc, S.W. Lowe, D.F. Tschaharganeh, and L.E. Dow. 2018. Optimized base editors enable efficient editing in cells, organoids and mice. Nat Biotechnol. 36:888–893.

Zimmermann, M., and T. de Lange. 2014. 53BP1: pro choice in DNA repair. Trends Cell Biol. 24:108–117.

